# β-catenin dependent axial patterning in Cnidaria and Bilateria uses similar regulatory logic

**DOI:** 10.1101/2020.09.08.287821

**Authors:** Tatiana Bagaeva, Andrew J. Aman, Thomas Graf, Isabell Niedermoser, Bob Zimmermann, Yulia Kraus, Magdalena Schatka, Adrien Demilly, Ulrich Technau, Grigory Genikhovich

## Abstract

In animals, body axis patterning is based on the concentration-dependent interpretation of graded morphogen signals, which enables correct positioning of the anatomical structures. The most ancient axis patterning system acting across animal phyla relies on β-catenin signaling, which directs gastrulation, and patterns the main body axis. However, within Bilateria, the patterning logic varies significantly between protostomes and deuterostomes. To deduce the ancestral principles of β-catenin dependent axial patterning, we investigated the oral-aboral axis patterning in the sea anemone *Nematostella* - a member of the bilaterian sister group Cnidaria. Here we elucidate the regulatory logic by which more orally expressed β-catenin targets repress more aborally expressed β- catenin targets, and progressively restrict the initially global, maternally provided aboral identity. Similar regulatory logic of β-catenin-dependent patterning in *Nematostella* and deuterostomes suggests a common evolutionary origin of these processes.

---

Graded morphogen signals comprise the top tier of the axial patterning cascades in Bilateria and their phylogenetic sister group Cnidaria (corals, sea anemones, jellyfish, hydroids) (*1-3*). Just like the posterior-anterior body axis of Bilateria, the oral-aboral body axis of Cnidaria is patterned by Wnt/β-catenin signaling (Fig. 1A) (*4, 5*). Although it is likely that β-catenin signaling is also involved in the axial patterning of earlier branching ctenophores and sponges (*6, 7*), cnidarians are the earliest branching animal phylum for which experimental gene function analyses are readily available. Thus, a cnidarian-bilaterian comparison can inform us about the ancient logic of the β-catenin-dependent axial patterning and mechanisms of molecular boundary formation. In this paper, we focus on the deciphering the mechanism of the oral-aboral axis patterning in the ectoderm of the early embryo of the sea anemone *Nematostella vectensis*.

**Figure 1.**
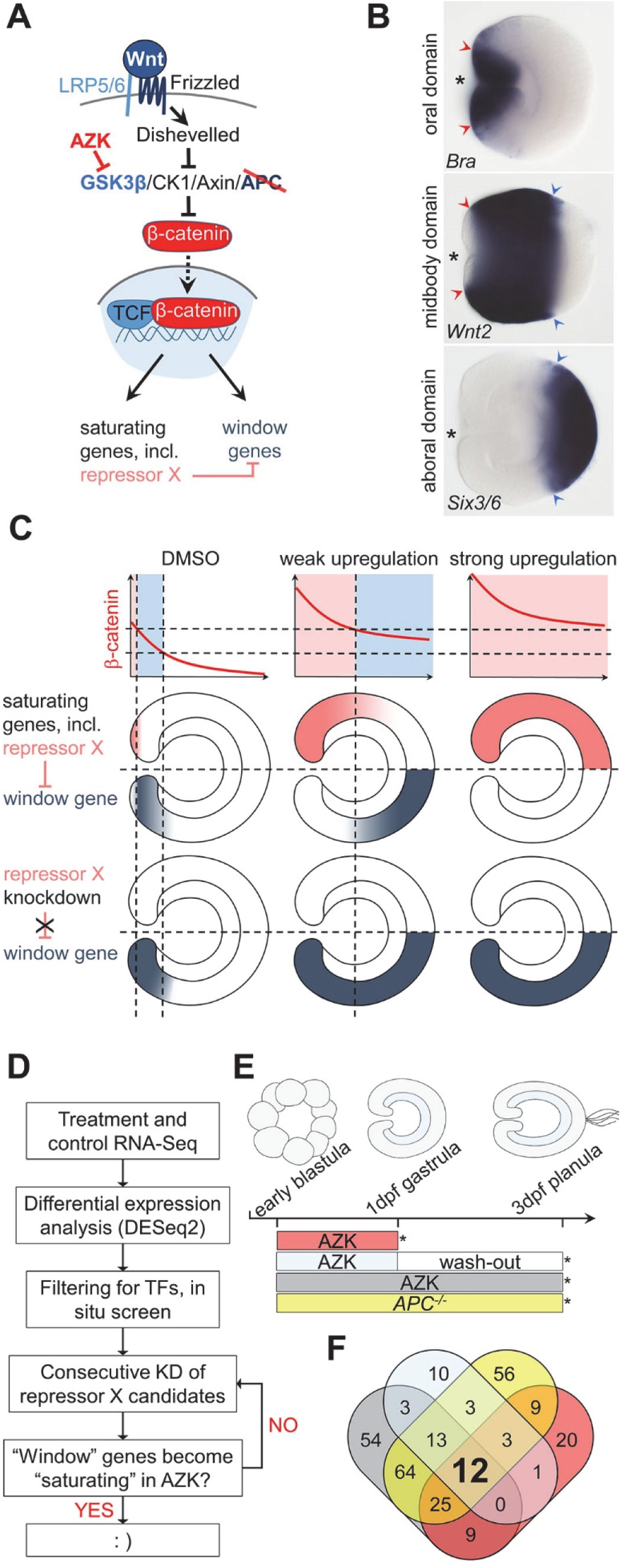
The “repressor X” concept and the search strategy. **(A)** Scheme of the Wnt/β-catenin signaling pathway indicating the members affected in this study in order to artificially upregulate it. We use two types of treatments (red) to upregulate β-catenin signaling: pharmacological inhibition of GSK3β by AZK and mutation of APC. **(B)** Oral, midbody, and aboral domains of the 1dpf gastrula visualized by molecular markers. Lateral views, oral to the left. Asterisk denotes the blastopore. Colored arrowheads demarcate corresponding positions. **(C)** Differential response of the “saturating” and “window” genes to different intensities of the β- catenin signaling and the hypothetical role of the transcriptional repressor X in regulating the “window” expression behavior. **(D)** Search strategy used to identify transcriptional repressor X. **(E)** Scheme of treatments. At 1dpf, AZK treatments were stopped at 24 hours post fertilization (hpf), and either RNA was extracted immediately, or the embryos were washed out and incubated in *Nematostella* medium until 3dpf (72 hpf). Asterisks indicate time points of RNA extraction. **(F)** Venn diagram with the numbers of the putative transcription factor coding genes upregulated by different treatments. The color code corresponds to that on (E).

Morphologically, the oral-aboral (O-A) axis in *Nematostella* becomes apparent at the onset of gastrulation, when future endoderm starts to invaginate, eventually forming the inner layer of this diploblastic organism. The establishment of the oral-aboral axis in *Nematostella* depends on β-catenin, which localizes to the nuclei at the future oral pole during early cleavage (*8*). Knockdown of β-catenin abolishes the O-A axis both morphologically and molecularly: the embryos fail to gastrulate and do not express oral ectoderm markers (*9*). In contrast, mosaic stabilization of β-catenin results in the formation of numerous ectopic oral structures or even complete ectopic axes (*4*) (see also fig. S1). By late gastrula stage, the ectoderm of *Nematostella* can be roughly subdivided into three axial domains: the oral domain characterized by *Brachyury* (*Bra*) expression, the midbody domain where *Wnt2* is expressed, and the aboral domain expressing *Six3/6* (Fig. 1B), whereas endodermal O-A patterning begins later in development (*10*). Pharmacological experiments, in which β-catenin signaling was upregulated by a range of concentrations of the GSK3β inhibitor 1-azakenpaullone (AZK) (Fig. 1A), showed that ectodermally expressed β-catenin dependent genes react to different levels of upregulation of β- catenin signaling dose-dependently and in two distinct ways (Fig. 1C) (*4*). Some genes, whose expression is normally restricted to the oral domain, increase their expression to saturation upon upregulation of β-catenin signaling and start to be expressed along the whole O-A axis at high AZK concentrations. We call them “saturating” genes below. Other genes, whose normal expression can be observed either in the oral domain or further aborally, require permissive “windows” of β-catenin signaling intensities. Upon weak pharmacological upregulation of β- catenin signaling, “window” gene expression shifts aborally, i.e. into the area where endogenous β-catenin signaling intensity is expected to be lower, while upon strong upregulation of β-catenin signaling their expression ceases altogether (*4*). A similar dose-dependent response to “windows” of β-catenin signaling intensity was previously demonstrated in axial patterning of deuterostomes (*11-14*), and possibly also in protostomes (*15, 16*). This similarity suggests that the regulatory principle behind the “window” behavior may represent the ancestral logic of β-catenin dependent axial patterning, however, its mechanism is not clear. Since this regulatory behavior is likely to be at the core of the O-A patterning in *Nematostella*, and possibly represent a general mechanism, we attempted to explain it. We hypothesized that both “saturating” and “window” genes are positively regulated by β-catenin (Fig. 1A). However, in order to account for the repression of the “window” genes upon upregulation of β-catenin, we postulated that there exists a “transcriptional repressor X”, which, being a “saturating” gene, becomes upregulated upon increased β-catenin signaling and inhibits the expression of the “window” genes (Fig. 1A, C). We set out to test our assumption and search for this hypothetical repressor.

Our hypothesis predicted that: i) the transcriptional repressor X is to be found among the “saturating” genes upregulated upon increased β-catenin signaling, ii) it has to be expressed in a contiguous domain along the O-A axis rather than in a salt-and-pepper manner to be able to act cell-autonomously, and that iii) the loss of function of the transcriptional repressor X will abrogate the β-catenin dependent repression of “window” genes converting them into “saturating” genes upon pharmacological upregulation of the β-catenin signaling (Fig. 1C). To test these predictions, we devised an RNASeq-based strategy for finding all transcription factors fulfilling these criteria (Fig. 1D). In order to obtain an off-target free list of transcription factors upregulated by β-catenin, we used two independent means of upregulating β-catenin signaling by suppressing the activity of two different members of the β-catenin destruction complex, which we further refer to as “treatments” (Fig 1A). First, we used AZK treatment spanning different time windows to suppress GSK3β. Second, we generated a line carrying a frameshift mutation in the *APC* gene (fig. S1). At 3 days post fertilization (3 dpf), all *APC*^*-/-*^ embryos display a phenotype similar to that of embryos incubated from early blastula on in AZK (fig. S1). Visual detection of the homozygous *APC* mutants at 1dpf is impossible, since the phenotype only becomes apparent at 2-3 dpf. However, an earlier study showed that “window” behavior of *Wnt2* persisted until at least 3 dpf (*5*), which suggested that the putative repressor X was expressed both at 1 dpf and at 3 dpf. Therefore, we compared the transcriptomes of 1 dpf embryos and 3 dpf embryos incubated in AZK with the transcriptomes of the 3dpf *APC*^*-/-*^ embryos (Fig. 1E; fig. S2), and controls. We then identified all putative transcription factor-coding genes upregulated by elevated β-catenin in all treatments by comparing our lists of differentially expressed genes with the list of gene models with a predicted DNA binding domain. We found twelve such putative transcription factors (Fig. 1F) of which we excluded five: two as metabolic enzymes falsely annotated by INTERPROSCAN (*17*) as transcription factors (*NVE21786* and *NVE12602*; for gene models see https://figshare.com/articles/Nematostella_vectensis_transcriptome_and_gene_models_v2_0/807696), one, *MsxC*, since it was not expressed in the wild type gastrula, and two, *Unc4* and *AshC* because they were expressed in single cells rather than in contiguous domains (fig. S3). The remaining seven candidates, *Bra, FoxA, FoxB, LIM homeobo*x (*Lmx), Shavenbaby (Svb), Dachshund* (*Dac*) and a putative Zn finger transcription factor *NVE11868*, were expressed in distinct continuous domains and displayed a typical “saturating” phenotype (fig. S3). In order to find out whether any these transcription factors were capable of repressing window genes, we individually knocked them down using shRNAs (fig. S4), incubated the knockdown embryos either in AZK or in DMSO and compared the expression of two well-characterized “window” genes *Wnt1* and *Wnt2* (*4*) in the knockdowns at late gastrula stage. Knockdowns of *Svb, Dac* and *NVE11868* led to no significant change in the expression of *Wnt1* and *Wnt2* in comparison to control shRNA (fig. S4). Therefore, these genes were also excluded from further analyses, and we focused on the remaining four candidates, *Bra, FoxA, FoxB*, and *Lmx*, and characterized their mutual expression domains and the effect of their knockdowns upon normal and pharmacologically enhanced β-catenin signaling (Suppl. Text, fig. S5-12).

The area of strong *Bra* expression overlaps with the *Wnt1* expression domain and abuts the *Wnt2* expression domain (fig. S5). Upon *Bra* knockdown, *Wnt1* expression was abolished not only in the AZK treatment but also in the DMSO treated controls, suggesting that *Wnt1* is positively regulated by *Bra* (Fig. 2, fig. S6). In contrast, *Wnt2* expression domain expanded orally in the DMSO controls and became ubiquitous upon the AZK treatment (Fig. 2). This suggests that Brachyury acts as the hypothetical transcriptional repressor X for *Wnt2*, but not for *Wnt1. FoxA* is expressed in the future pharynx of the embryo and in the domain immediately around the blastopore inside the ring of *Wnt1* expressing cells (fig. S5). *FoxA* knockdown does not affect *Wnt2* expression, but *Wnt1* expression becomes stronger and expands further orally in DMSO and globally in the AZK treatment (Fig. 2). Thus, FoxA appears to act as the hypothetical repressor X for *Wnt1*, but not for *Wnt2. FoxB* is co-expressed with *Bra* in the domain where *Bra* expression is strong (fig. S5), i.e. abutting the *Wnt2* expression domain, and *Lmx* is a weakly expressed gene active in a domain starting from the *Wnt1* expressing cells and quickly fading out further aborally (fig. S5). *FoxB* knockdown resulted in the expansion of both *Wnt1* and *Wnt2* expression in AZK, but the staining appeared weak, and *Lmx* RNAi effect on *Wnt1* and *Wnt2* largely recapitulated the effect of *Bra* RNAi, albeit milder (Fig. 2). Single, double and triple knockdown experiments suggest that the role of these two transcription factors appears to be in supporting the activity of Bra and FoxA in the areas, where they are co-expressed (Suppl. Text, fig. S6-12). Simultaneous knockdown of *Bra, Lmx, FoxA* and *FoxB* with a mixture of shRNAs (shBLAB) completely abolishes the oral identity of the embryo at the molecular level: the midbody marker *Wnt2* shifts orally, expanding all the way to the bottom of the pharynx in DMSO, while the *Wnt2*-free aboral domain expands (Fig. 2). A much more pronounced expansion of the aboral domain and the confinement of the midbody marker *Wnt2* to the oralmost part of the embryo upon the combined knockdown of *Bra* together with either *Lmx* or *FoxB* or both in comparison to the individual *Bra* knockdown shows that the functions of these genes are non-redundant (Fig. 2; fig. S6-9). We conclude that oral “window” genes are activated by β- catenin signaling (either directly or indirectly), and repressed by β-catenin-dependent “saturating” transcription factors. No single transcriptional repressor X exists, but rather *Bra, Lmx, FoxA* and *FoxB* appear to be the unit defining oral identity in the *Nematostella* embryo. Strikingly, the knockdown of any of these four transcription factors did not prevent the normal gastrulation, and all the effects at this developmental stage remained purely molecular, pointing at the potential role of maternal factors in the gastrulation process (Suppl. Text., fig. S13).

**Figure 2.**
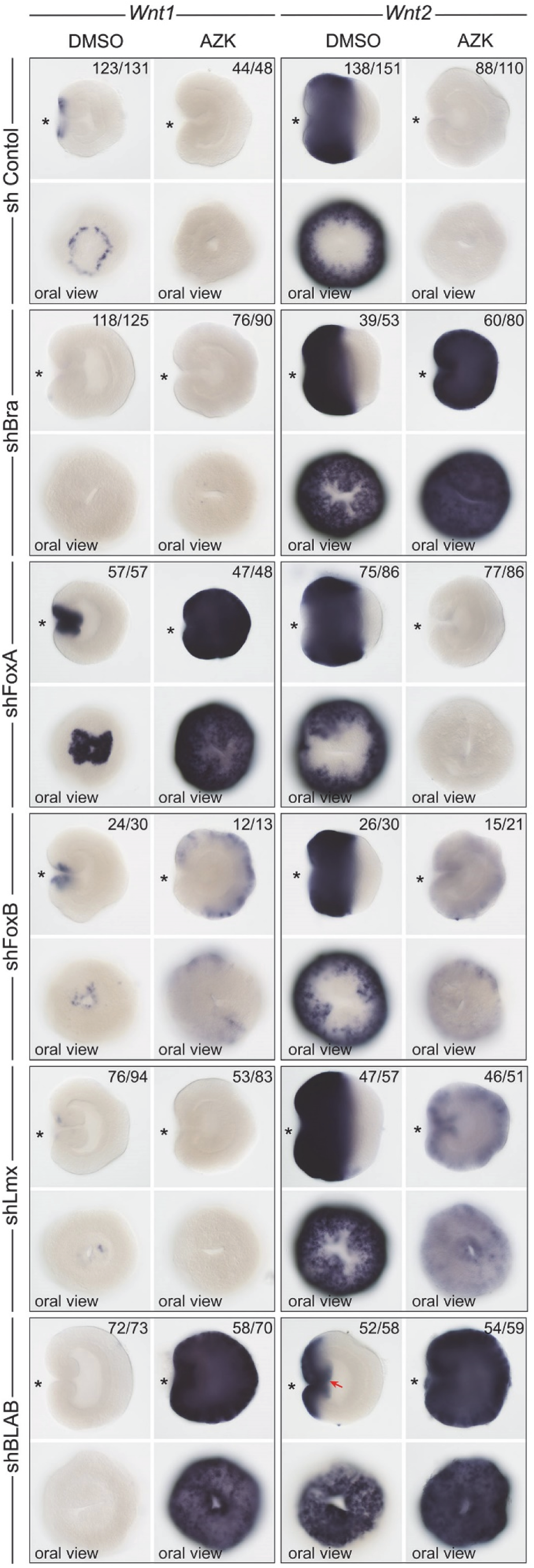
The effect of the repressor X candidates knockdown on the expression of the “window” genes Wnt1 and Wnt2. *Bra* and *Lmx* knockdowns convert *Wnt2* into a saturating gene, while *FoxA* knockdown does the same with *Wnt1*. The effect of *Lmx* knockdown appears to be similar but weaker than that of *Bra. FoxB* knockdown results in a “weak AZK effect” on both *Wnt1* and *Wnt2* suggesting that FoxB mildly represses both. The effects of the knockdowns of *Bra, Lmx*, and *FoxB* on *Wnt2* expression are non-redundant, but similar and additive (see fig. S8, 9). Quadruple knockdown with shRNA against *Bra, Lmx, FoxA* and *FoxB* (=shBLAB) removes oral molecular identity of the embryo completely. Red arrow indicates the bottom of the pharynx expressing the midbody marker *Wnt2*. On lateral views, asterisk denotes the blastopore.

Previous work demonstrated that the aboral markers *FoxQ2a* and *Six3/6*, which are downregulated upon elevated β-catenin signaling, still require some β-catenin signaling in order to be expressed (*9*). Therefore, it is conceivable that the patterning logic we discovered for the oral domain may be applicable to the whole of the O-A patterning, with more orally expressed β- catenin-dependent genes acting as transcriptional repressors for the more aborally expressed β- catenin-dependent genes. To test that, we investigated the mechanism of the maintenance of the other clear molecular boundary present in late gastrula ectoderm: the one between the *Wnt2*- positive midbody domain and the *Six3/6*-positive aboral domain (Fig. 1B, fig. S5). If the proposed regulatory logic were correct, there would have to be at least one “transcriptional repressor Y”, which: i) has to be expressed in the midbody domain, ii) has to be positively regulated by β-catenin and repressed by the oral, “saturating” transcription factors (i.e. it has to be encoded by a “window” gene), and iii) has to counteract the oral expansion of the aboral domain. Since “window” genes are downregulated upon elevated β-catenin signaling, we looked at the transcription factor coding genes downregulated by all treatments in our RNASeq experiment (Fig. 3A). We found 25 such genes, and two of them attracted our attention as members of the Sp family of the Krüppel-like transcription factors, which are important regulators of axial patterning and mediators of β-catenin-signaling in Bilateria (*18-23*) and in *Hydra* (*24*). *Nematostella* has representatives of all three Sp families present in Bilateria. Like in chordates (*20, 23*) and in *Hydra* (*24*), *Nematostella Sp5* is activated by β-catenin signaling. It is a “saturating” gene expressed in the domain similar to that of *Axin-like* and *Tcf* (fig. S1, fig. S14). In contrast, *Sp1-4* and *Sp6/9* are expressed in the aboral and the midbody domain respectively (fig. S14; see also (*25*)), and are among the genes downregulated in all treatments (Fig. 3A). In addition to the broad expression in the midbody domain (bordering the *Bra* domain orally and the *Six3/6* domain aborally; fig. S5), *Sp6-9* is also strongly expressed in individual cells scattered all over the embryo. This single cell expression is not dependent on β-catenin signaling (fig. S14). Since the expression of *Sp6-9* in the midbody domain suggested it as a candidate for the role of the transcriptional repressor Y, we tested whether it fulfilled the two remaining transcriptional repressor Y criteria set above. We could show that the knockdown of the four oral transcription factors *Bra, FoxA, FoxB* and *Lmx* expanded the expression of *Sp6-9* orally in DMSO and globally in AZK (Fig. 3B), i.e. *Sp6-9* behaved as a “window” gene. Moreover, the knockdown of *Sp6-9* resulted in the oral expansion of *Six3/6* (Fig. 3C). Predictably, since *Bra* knockdown results in the oral shift of the midbody domain (Fig. 2, Fig. 3B) and expansion of the aboral domain (Fig. 3C), *Six3/6* expansion was much more pronounced upon the combined knockdown of *Sp6-9* and *Bra* (Fig. 3C). Thus, Sp6-9 appears to act as hypothetical repressor Y at least for *Six3/6*, which suggests that the regulatory logic we proposed is applicable not just to the oral domain but to the whole β-catenin-dependent oral-aboral axis patterning in *Nematostella*.

**Figure 3.**
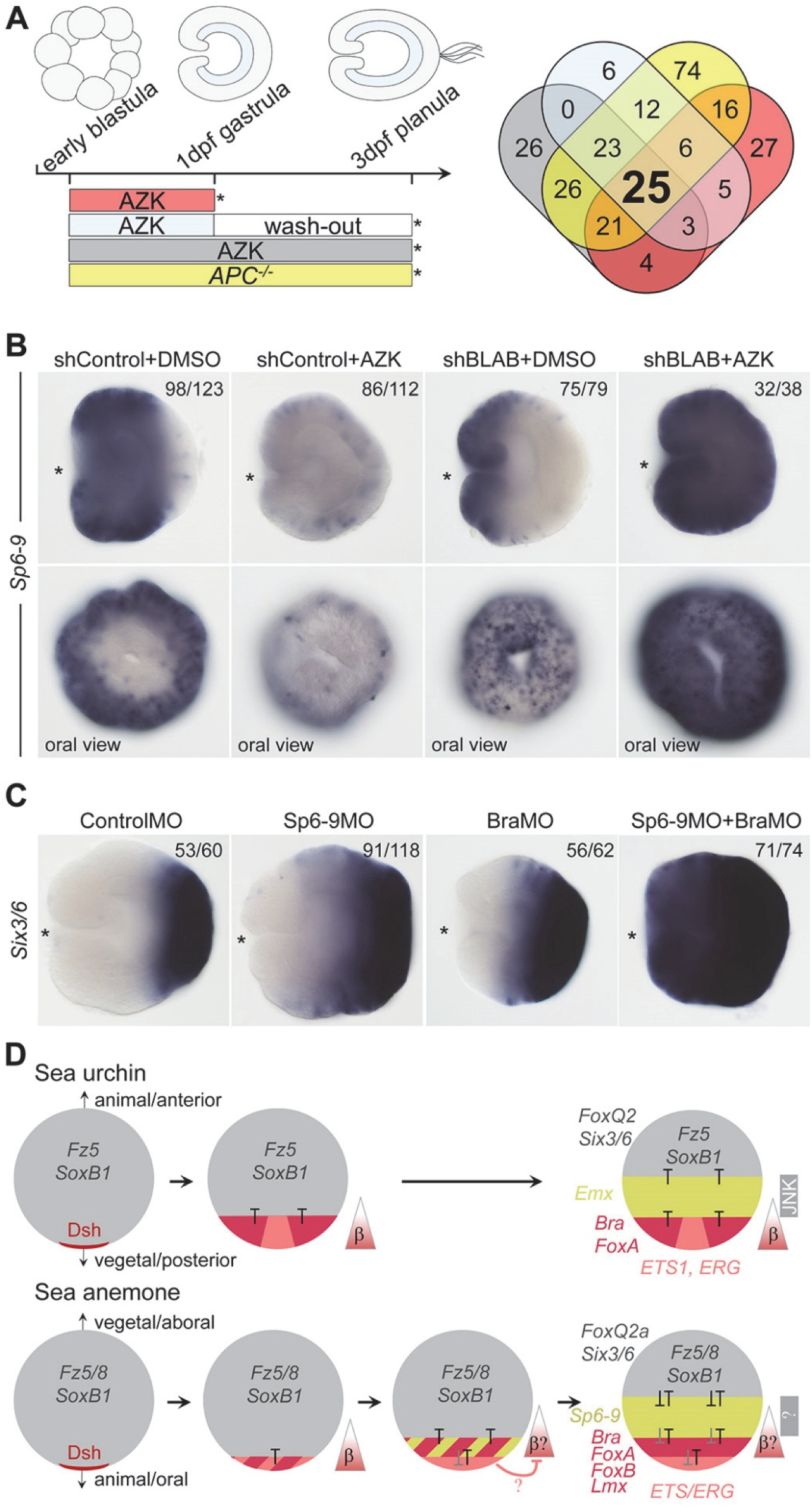
Midbody domain prevents oral expansion of the aboral domain. **(A)** Scheme of the treatments (as in Fig. 1) and Venn diagram showing the number of putative transcription factors downregulated by various treatments. **(B)** *Sp6-9* is a “window” gene shifting orally upon simultaneous knockdown of *Bra, Lmx, FoxA* and *FoxB* (=shBLAB) and expanding globally upon shBLAB knockdown followed by AZK treatment. Sp6-9-free area disappears in shBLAB. Lateral views, oral to the left; asterisk denotes the blastopore. **(C)** Sp6-9 prevents oral expansion of the aboral marker *Six3/6*. In BraMO, *Six3/6* expression is also expanded orally, likely due to the oral shift of the *Sp6-9* expression (see fig. S7). Oral expansion of *Six3/6* is enhanced upon double knockdown of *Sp6-9* and *Bra*. Lateral views, oral to the left; asterisk denotes the blastopore. **(D)** Comparison of the early β-catenin dependent patterning in sea urchin and *Nematostella* shows clear similarities. Unfertilized egg with maternal *Fz5/8* and *SoxB1* mRNA (future anterior/aboral markers) and maternal Dsh protein localized at the gastrulation pole (*38, 39*). Upon activation of β-catenin signaling in the embryo, first in the endomesodermal domain and then in the posterior/oral ectoderm the expression of *Fz5/8* and *SoxB1* is suppressed, and the anterior/aboral markers (including the zygotic genes *Six3/6* and *FoxQ2*) become progressively confined to one side of the axis. The axis becomes patterned by mutually repressive transcription factors (T). Grey “T” in *Nematostella* indicate repressive interactions, for which candidate transcription factors are not known. Triangles with a β denote the direction of the β-catenin signaling gradient. β? indicates that in *Nematostella*, nuclear β-catenin could only be experimentally detected until midblastula stage (*9*), after which the presence of nuclear β-catenin gradient is deduced based on target gene response. After preendodermal plate is specified in *Nematostella*, β-catenin signaling becomes repressed there by an unknown mechanism (*9*), possibly involving ERG (*25*).

We demonstrated that the logic of the β-catenin dependent O-A patterning relied on more orally expressed β-catenin targets displacing the expression domain of the more aborally expressed β-catenin targets further aborally. Therefore, we decided to test whether aboral fate represented the default state of the whole *Nematostella* embryo, which then became progressively restricted to the aboral domain by the orally expressed β-catenin-dependent factors, as it is described for the anterior ectodermal domain in deuterostomes (*12, 26-28*). The fact that the major aboral determinant *Six3/6* requires an initial β-catenin signal in order to be expressed (*9*) argues against this otherwise very appealing hypothesis. However, *Six3/6* is a zygotic gene, whose expression becomes detectable at 12 hours post fertilization (hpf), which is 4 hours later than the onset of expression of the oral marker *Bra* (fig. S15). Notably, even the earliest expression of *Six3/6* is not ubiquitous, but localized to the future aboral side of the O-A axis. However, we do find aboral markers, whose expression is initially maternal and ubiquitous and subsequently becomes restricted to the aboral end in a β-catenin-dependent manner (fig. S15). One of them is *Frizzled 5/8* (fig. S15), which was shown to be a negative regulator of *Six3/6* and *FoxQ2a* in *Nematostella* and sea urchin (*9, 27, 29*). The other one is *SoxB1* (fig. S15), whose initially ubiquitous expression is cleared out of the organizer and endomesodermal area in deuterostomes (*30, 31*). Individual or simultaneous knockdowns of the oral and midbody factors Bra and Sp6-9 in *Nematostella* significantly expand the expression domain of *SoxB1* (fig. S16). Although qPCR data suggest that sea urchin SoxB1 is a positive maternal upstream regulator of *FoxQ2* (*32*), the negative effect of *SoxB1* knockdown on *Six3/6* and *FoxQ2a* expression in *Nematostella* is not pronounced (fig. S16), and it is still unclear what kind of positive regulatory input maintains the aboral expression of *Six3/6* and hence other aboral markers. Nevertheless, our data clearly support the aboral-by-default model.

In summary, here we showed that *Bra, FoxA, Lmx* and *FoxB* define the oral molecular identity of the *Nematostella* embryo and prevent oral expansion of the more aborally expressed β-catenin targets. We also identified *Sp6-9*, a “window” gene expressed in the midbody domain, as the agent preventing the oral expansion of the aboral domain. The whole *Nematostella* embryo initially represents an aboral ectodermal territory, which is established maternally. During the first day of development, this territory becomes restricted to the aboral end of the O-A axis in a β-catenin-dependent manner by “saturating” and “window” transcriptional repressors, which form mutually repressive pairs capable of generating sharp domain boundaries (Fig. 3D). Comparison with sea urchin reveals remarkable conservation of the components of the axial patterning gene regulatory network downstream of β-catenin. Also in sea urchin, *Bra* and *FoxA* are central in the β-catenin dependent axial patterning of the blastoporal domain (*33, 34*). The midbody domain of the sea urchin embryo appears to be defined by an Antennapedia class transcription factor Emx (*35, 36*), rather than by a Krüppel-like factor Sp6-9. However, the patterning of the apical ectoderm is again accomplished by the same components in the sea anemone and sea urchin embryos (*27, 37*). Importantly, not only the genes involved, but also the regulatory logic of gradual restriction of the apical ectodermal territory by β-catenin-dependent transcription factors appear to be highly similar in the β-catenin dependent oral-aboral patterning in the anthozoan *Nematostella* and in the posterior-anterior patterning in sea urchin and other investigated deuterostomes, including vertebrates (Fig. 3D) (*11-13, 27, 28, 33, 34*). We conclude that these processes share a common evolutionary origin predating the cnidarian-bilaterian split.

## Acknowledgements

We are grateful to the Core Facility for Cell Imaging and Ultrastructure Research of the University of Vienna for the access to the confocal and scanning electron microscopes, Patrick Steinmetz for the fluorescent ISH protocol, Patricio Ferrer Murguia for an shRNA against *FoxA*, Saskia Hartmann for the initial genotyping of the APC mutants, Rohit Dnyansagar for the help with bioinformatics, Emmanuel Haillot for discussions, and David Mörsdorf for critically reading the manuscript.

## Author contributions

T.B. performed the majority of the experiments, planned experiments and analyzed data; A.A and U.T. conceived the generation of the *APC* mutant line; A.J.A. generated the *APC* mutant line, and started its characterization together with A.D.; T.G. and I.N. performed treatments, and prepared RNA for sequencing; B.Z. supervised the bioinformatic analysis; Y.K. performed transplantations on *Bra* morphants; M.S. generated mosaic *EF1α::β-cat_stab* polyps; G.G. conceived the study, planned experiments, performed experiments, analyzed data and wrote the paper. All authors edited the paper.

## Data and materials availability

All data needed to evaluate the conclusions in the paper are present in the paper or the supplementary materials. Raw RNASeq reads are accessible at NCBI (BioProject PRJNA661731).

## Supplementary materials

Materials and methods

Supplementary text

Figures S1-S16

Tables S1 and S2.

References 40-51

