## Supplementary information for "β-catenin dependent axial patterning in Cnidaria and Bilateria uses similar regulatory logic"

**Contents:**

Materials and methods

Supplementary text

Figures S1-S16

Tables S1 and S2.

References 40-51

### Materials and methods

#### Animal husbandry, microinjection and *APC* mutant line

*Nematostella* polyps were kept in *Nematostella* medium (16‰ artificial seawater, NM) at 18° in the dark and induced to spawn by placing into a 25°, illuminated incubator for 10 hours. The eggs were fertilized for 30 minutes and dejellied in 3% L-cysteine/NM and washed 6 times in NM. Microinjection was performed under the Nikon TS100F microscope using Eppendorf Femtojet and Narishige micromanipulators. The *APC* mutant line was generated by injecting *Nematostella* zygotes with 500 ng/µl single gRNAs (protospacer 5'CACAGCTATGAGGGCCAC) and 500 ng/µl nls-Cas9 (PNA Bio, Thousand Oaks, CA, USA). Mosaic F0 animals were crossed to produce $APC^{+/-}$ F1 carrying a single T insertion after the position 331 of the coding sequence of *Nematostella APC* (Genbank KT381584). Heterozygous F1 were crossed to obtain F2. In situ hybridization analysis showed that 27% of the F2 embryos expressed *Bra* throughout the ectoderm of the gastrula, while 73% had normal *Bra* expression (N=221). At 3dpf, 10 out of 10 randomly selected F2 embryos demonstrating the typical bagel phenotype similar to that of the AZK treated embryos proved to be $APC^{-/-}$ when genotyped by Sanger sequencing of the mutated locus (fig. S1F). For genotyping live polyps, individual primary polyps or tentacle clips were fixed by 3 washes in 100% methanol, aspirated, and dried for 20 min at 50°C with the tube lids open. Then, samples were digested in 30 µl of extraction buffer (10mM Tris-HCl pH8, 1 mM EDTA pH8, 25 mM NaCl, 200 µg/ml proteinase K) for 2 h at 50°C, and proteinase K was inactivated by heating the samples to 95°C for 5 min. After proteinase K inactivation, 3 µl of the digest was used as template for the PCR with the primers flanking the locus recognized by the gRNA (APCspF 5'AGAATCCTGCAGAAGATGAACA, APCspR 5'CCTGGCATACAAAGGTGACA). The PCR product was purified and directly sequenced with the APCspF primer. For genotyping embryos after in situ hybridization, the embryos were dehydrated in ethanol series, washed twice with 100% ethanol, embedded into Murray's clear solution (benzyl benzoate:benzyl alcohol = 2:1), imaged, washed several times in methanol and then processed as described above.

#### Pharmacological treatments, gene knockdown and gene overexpression

5µM 1-azakenpaullone (AZK) used for the treatments was prepared by diluting 5 mM AZK dissolved in DMSO with NM. Equal volume of DMSO was used to treat the control embryos. The time windows of the treatments are presented in Fig. 1E. Gene knockdowns were performed by electroporation with shRNA as specified (*40, 41*). Two non-overlapping shRNAs were used for each of the genes to confirm the specificity of the observed phenotypes except for the case of *Brachyury* and *Sp6-9*, where two or one shRNAs and one translation-blocking morpholino (MO) were used (Tables S1-S2). shRNA against *mOrange* was used as a control for all other shRNA and a control MO we described previously (*4*) was used to control for the BraMO and Sp6-9MO phenotypes. The RNAi efficiency was estimated by Q-PCR, and the activity of the BraMO and Sp6-9MO was assayed by co-injecting it with the wild type and 5-mismatch mRNA containing the morpholino recognition sequences fused to mCherry (fig. S4). Capped mRNA was synthesized using mMessage mMachine kit (Life Technologies) and purified with the Monarch RNA clean-up kit (NEB). Bra and FoxB mRNA for overexpression was also produced as described above. A stabilized form of β-catenin was generated by removing the first 240 bp of the β-catenin coding sequence as described in (*42*). An ATG was added, and the fragment, which we called β-cat_stab, was cloned into an expression vector downstream of the ubiquitously active *EF1α* promoter (*43*). Mosaic expression of the *EF1α::β-cat_stab* was achieved by meganuclease-assisted transgenesis, as described (*44*). Primers against GAPDH were used as normalization control in QPCR.

**Transcriptome sequencing and analysis**

Total RNA was extracted with TRIZOL (Life Technologies) or with GeneElute Mammalian Total RNA Miniprep Kit (Sigma) according to the manufacturer's protocol; poly-A enriched mRNA library preparation (Lexogen), quality control, and multiplexed Illumina HiSeq2500 sequencing (50 bp, single-end) were performed at the Vienna BioCenter Core Facilities. The number of the sequenced biological replicates of different treatments is shown in fig. S2. The reads were aligned with STAR (*45*) to the *Nematostella vectensis* genome (*46*) using the ENCODE standard options, with the exception that --alignIntronMax was set to 100 kb. Hits to the gene models v2.0 (https://figshare.com/articles/Nematostella_vectensis_transcriptome_and_gene_models_v2_0/807696), were tallied with featureCounts (*47*), and differential expression analysis was performed with DeSeq2 (*48*). Expression changes in genes with Benjamini-Hochberg adjusted p-value < 0.05 were considered significant. No additional expression fold change cutoff was imposed. Transcription factor candidates were determined analyzing the transcriptome with INTERPROSCAN (*17*) and filtering for genes containing the domains described in (*49*). The intersection between the latter set and our differentially expressed genes comprised the models of putative transcription factors.

**In situ hybridization, fluorescent double in situ hybridization, SEM**

In situ hybridization was performed exactly as described in(*4*) with a minor change in the fixation protocol: here, we fixed the embryos for 1 hour in 4% PFA/PBS at room temperature and washed the embryos several times first in PTw (1x PBS, 0.1% Tween 20) and then in 100% methanol prior to storing them at -20°C. Imaging was performed with a Nikon 80i compound microscope equipped with the Nikon DS-Fi1 camera. For the fluorescent double in situ hybridization, the hybridization protocol was similar to the single chromogenic in situ protocol except for the changes outlined below. FITC- and Dig-labelled RNA probes were simultaneously added to the sample. After stringent post-hybridization washes, the embryos were blocked in the 0.5% TSA Blocking Reagent (Perkin-Elmer) in TNT buffer for 1 h, and stained overnight at 4°C with anti-Dig-POD antibody (Roche) diluted 1:100 in blocking buffer. The unbound antibody was then removed by 10x10 min TNT washes, and the fluorescent signal was developed using the TSA Plus Cyanine 3 System (Perkin-Elmer) according to the manufacturer's protocol. The staining was stopped by multiple TNT washes, and peroxidase was inactivated by a 20 min wash in 1% $H_2O_2$/TNT in the dark. After that, the embryos were washed several times with TNT, blocked as described above and stained with the anti-Fluorescein-POD antibody (Roche) diluted 1:50 in blocking buffer. Fluorescent signal was then developed as described above using the TSA Plus Fluorescein System (Perkin-Elmer). After stopping the staining with multiple TNT washes embedded in Vectashield (Vectorlabs) and imaged with the Leica SP8 CLSM. Preparation of the samples for the SEM was performed as described in (*4*). Imaging was done using the JEOL IT 300 scanning electron microscope.

**Supplementary text:**

**Topology of the gene regulatory network**

In order to understand the genetic interactions between the four factor X candidates, we analyzed their expression in individual, double, triple and quadruple knockdowns (fig. S7-12). While identification of the exact topology of this network requires stage-by-stage ChIP data for all the transcription factors from the onset of their expression until late gastrula, we can suggest a possible topology based on genetic interactions. Upon shRNA mediated knockdown of *Bra*, *FoxB* and *Lmx* become abolished. *FoxA* expression is confined to the bottom of the forming pharynx, where *Bra* is not expressed, suggesting that Bra activates the expression of these three genes in the area where they are normally co-expressed with *Bra* (fig. S7). The effect of *Bra* knockdown on *Bra* expression is more complex: while shRNA-mediated knockdown reduces the amount of *Bra* mRNA (fig. S7), the translation blocking morpholino mediated knockdown of *Bra* upregulates the expression of *Bra* gene (fig. S10A). The most likely explanation for this is that Bra protein may act as a transcriptional repressor of the *Bra* gene. *Bra* knockdown also abolishes *Wnt1* and *Wnt3* expression in DMSO and in AZK, however *Wnt1* is expressed in AZK in quadruple knockdowns (Fig. 2, fig. S13) and in triple knockdowns, when shFoxA is used (fig. S9). Thus, the regulation of *Wnt1* by β-catenin may still be direct. *Wnt3*, in contrast, is abolished in AZK in quadruple knockdowns. This suggests that *Wnt3* is either regulated by β-catenin indirectly via Bra and FoxB (fig. S13, and see below), or, more likely, that a yet unidentified "window" transcriptional repressor preventing aboral expansion of the *FoxA*/*Wnt1*/*Wnt3* boundary (fig. S5) becomes de-repressed in AZK upon quadruple knockdown and suppresses *Wnt3* (just like *Wnt2* becomes de-repressed in shBLAB and expands orally).

According to double and triple knockdowns followed by AZK treatment, *Bra, FoxB*, *Lmx* and *FoxA* are likely to be direct β-catenin targets (fig. S8, 9). Upon individual *FoxB* shRNA knockdown, in DMSO, the ring of *Wnt1* concentrates around the blastopore opening and appears weaker, and in AZK, *Wnt1* expression expands aborally but remains weak and absent in blastopore lips (Fig. 2). *Wnt2* expression in *FoxB* knockdown is slightly expanded orally in DMSO and globally in AZK, but also remains weak (Fig. 2). This is strikingly similar to the effects we documented in wild type embryos treated with lower concentrations of AZK(*4*). The effect of its knockdown on the expression of the *Wnt* genes suggests that FoxB might act as an enhancer of Bra and FoxA activity. The role of *Lmx* is also not fully clear, since the effect of its knockdown on *Wnt1* and *Wnt2* seems to be weaker but similar to the effect of the *Bra* knockdown (Fig. 2), suggesting that Bra and Lmx proteins might cooperate in regulating the same targets. The only observable difference is that *Lmx* is not co-expressed with *Wnt3* (fig. S5), which is localized to the *FoxA/Bra* co-expression domain, and *Lmx* knockdown does not seem to strongly affect *Wnt3* expression (fig. S13). The role of Lmx and FoxB appears to be in supporting the expression and function of *Bra* in their respective co-expression domains (fig. S5, 7-9, see also main text), while strong FoxA appears to suppress *Bra* expression in the absence of FoxB (i.e. at the bottom of the pharynx, fig. S7). Similar expression of *Bra* upon single knockdown of *FoxB* (fig. S7), double knockdown of *FoxB*+*Lmx* or *FoxB*+*FoxA* (fig. S8), and triple knockdown of *FoxB*+*FoxA*+*Lmx* (fig. S9) points towards the critical role of FoxB in maintaining the normal domain of strong *Bra* expression. One yet unclear effect is the nearly normal expression of *Bra* (normal domain plus bottom of the pharynx) and upregulation of *FoxB* upon double knockdown of *Lmx* and *FoxA* (fig. S8), although single knockdowns of these genes reduced *Bra* expression restricting it to the bottom of the pharynx (fig. S7). In spite of the effects of the *FoxB* knockdown and the *Bra* knockdown on *FoxA* expression being highly similar (expression is confined to the bottom of the pharynx, fig. S7), overexpression of *FoxB* in the shBra background does not rescue the *FoxA* phenotype (fig. S11). In contrast, overexpression of *Bra* mRNA in the shFoxB background completely rescues the shFoxB effect on *FoxA* without inducing *FoxA* expression outside of its normal domain (fig. S11). The lack of ectopic expression of *FoxA* upon *Bra* overexpression suggests that the genes encoding transcription factors repressing *FoxA* aborally are not under Bra control. Identical effects of *Bra*, *FoxB* and *Lmx* knockdowns on the expression of the midbody marker *Sp6-9* (fig. S7) suggests that these factors might co-operate in preventing the oral expansion of the expression domain of this gene. However, Bra is clearly the key player in this inhibition: co-injection of shBra with *FoxB* mRNA does not fully suppress *Sp6-9* expression, although *Sp6-9* appears to be weaker than in control, and its oral expansion appears to be less pronounced in comparison to the shBra alone (compare fig. S7 and 11). In contrast, co-injection of shFoxB with *Bra* mRNA drastically reduces *Sp6-9* expression (fig. S11). Notably, *Sp6-9* knockdown does not cause aboral expansion of *Bra*, *FoxA*, *FoxB* and *Lmx* (fig. S10B), suggesting that they are suppressed by the yet unidentified midbody genes. Similarly, it is unclear what prevents aboral expansion of the other oral markers expressed in concentric rings, e.g. Wnt1, Wnt3, Wnt4 and WntA (see fig. S5). In contrast, Sp6-9 clearly prevents oral expansion of the aborally expressed *Six3/6* and *Sp1-4* (Fig. 3C; fig. S10B,C). Conversely, *Six3/6* knockdown results in the aboral expansion of *Sp6-9* (although the penetrance is not too high at 55%, N=55) (fig. S10D). Since the suppression of *Sp6-9* alone does not result in the aboral expansion of *Bra*, it is clear that some yet unidentified factors are involved in preventing aboral expansion of the oral and midbody genes. The deduced topology of the genetic interactions in this GRN is presented in the fig. S12.

**Knockdown of the four oral TFs affects de novo axis formation but not the normal gastrulation**

Our analysis of the effects of the knockdowns of the four transcription factors defining oral identity in *Nematostella* provided another highly surprising result. Although gastrulation in *Nematostella* is abolished by β-catenin morpholino (*9*), and ectodermal co-expression of *Wnt1* and *Wnt3* is sufficient to induce axis and germ layer formation at any positon in the embryo (*4, 50*), neither individual knockdowns nor the quadruple knockdown of *Bra*, *Lmx*, *FoxA* and *FoxB*, affected the process of gastrulation in any detectable way. This was unexpected, since *Bra* knockdown abolishes both *Wnt1* and *Wnt3*, and *Lmx* knockdown abolishes *Wnt1* (Fig. 2; fig. S6, 7, 10, 13). In order to address this discrepancy, we tested whether the expression of our four candidate transcription factors affects ectopic axis induction by blastopore lip transplantation (fig. S13). We predicted that if any of these molecules were necessary for axis induction, their loss from the donor blastopore lip tissue would abolish its inductive potential. Indeed, the knockdowns of *Bra* and *Lmx* nearly abolished the inductive capacity of the blastopore lip when compared to the transplantations from Control MO or shControl injected embryos (Z-test, $p<1e-5$ in both cases), and *FoxB* knockdown significantly reduced it (Z-test, $p<0.01$). This latter effect might be due to the reduced *Wnt1* and *Wnt3* expression in shFoxB embryos (Fig. 2, fig. S13). In contrast, the knockdown of *FoxA*, which appears to be the transcriptional repressor of *Wnt1*, strongly increases induction efficiency (Z-test, $p<1e-5$). This suggests that all these factors affect the ectopic axis and germ layer formation, while normal gastrulation in the embryo relies primarily on maternally deposited determinants.

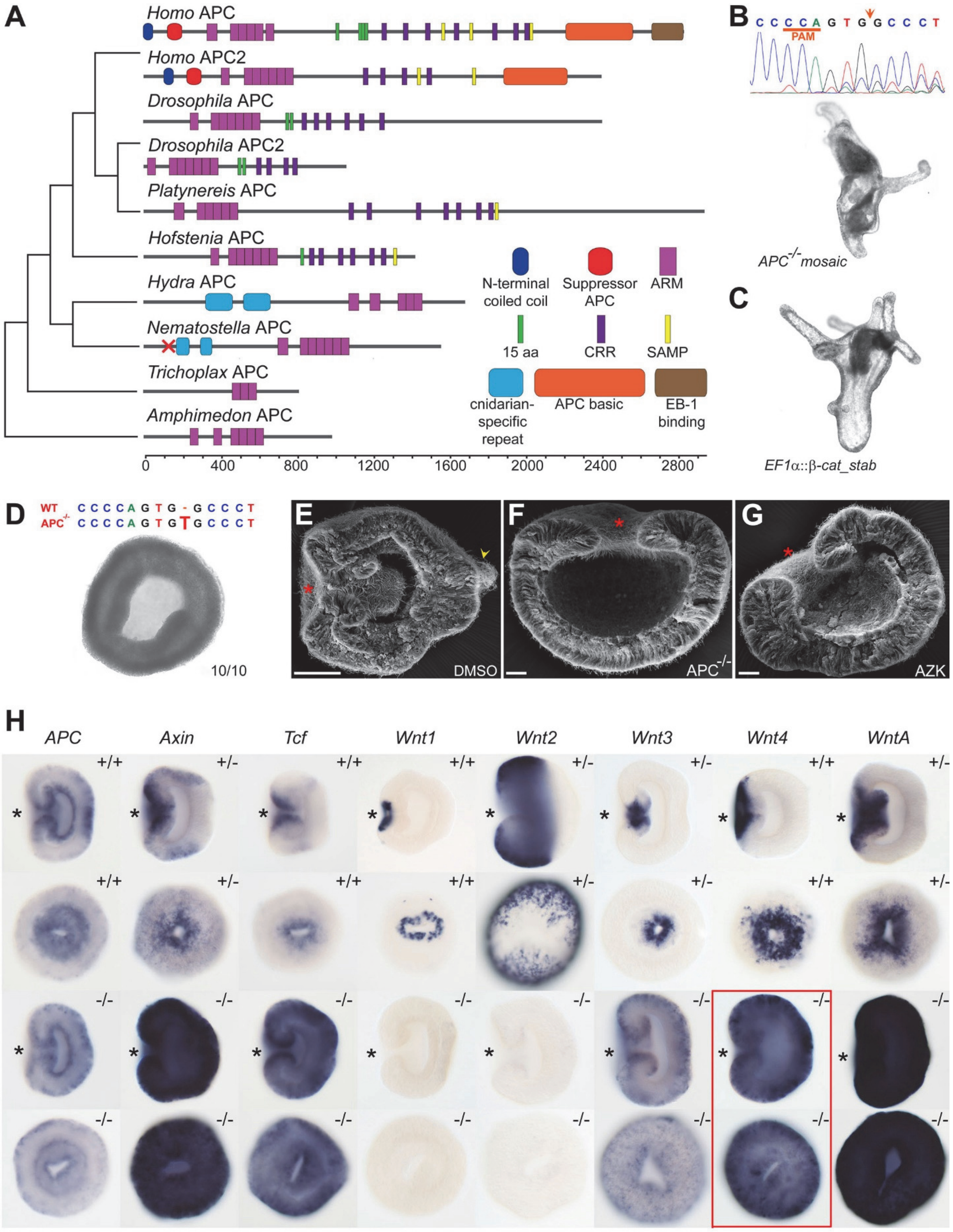
A
Homo APC
Homo APC2
Drosophila APC
Drosophila APC2
Platynereis APC
Hofstenia APC
Hydra APC
Nematostella APC
Trichoplax APC
Amphimedon APC
N-terminal coiled coil
Suppressor APC
ARM
15 aa
CRR
SAMP
cnidarian-specific repeat
APC basic
EB-1 binding
0
400
800
1200
1600
2000
2400
2800
B
C C C C A G T G G C C C T
PAM
APC-/- mosaic
C
EF1α::β-cat_stab
D
WT C C C C A G T G - G C C C T
APC-/- C C C C A G T G T G C C C T
10/10
E
DMSO
F
APC-/-
G
AZK
H
APC
Axin
Tcf
Wnt1
Wnt2
Wnt3
Wnt4
WntA
+/+
+/-
-/-

Fig. S1

**Supplementary Figure 1 Characterization of the *APC* mutants.**
(**A**) SMART annotation (*51*) of the domain structure of the animal APC proteins. Cnidarian APCs have armadillo repeats (ARM), but appear to be missing the typical 15 and 20 amino acid repeats (15 aa, CRR) present in Bilateria and used for β-catenin binding. SAMP – Axin binding domain. Red cross on the *Nematostella* protein indicates the position of the frameshift mutation.

(**B** to **C**) Although several important domains are missing in the non-bilaterian APC proteins, *Nematostella* APC appears to act via β-catenin. (B) Mosaic *APC* mutant develops multiple ectopic oral structures. Upon genotyping of this polyp, the sequencing chromatogram shows the accumulation of extra peaks around the Cas9 cutting site (arrow). (C) Mosaic expression of the stabilized form of the *Nematostella* β-catenin results in a comparable phenotype.

(**D**) Genotyping shows that 10/10 3 dpf F2 embryos demonstrating the oralization phenotype are homozygous *APC* mutants with a T insertion. The image shows a representative *APC* mutant (oral view).

(**E** to **G**) SEM image of a control 3 dpf planula (E) with an elongated oral-aboral axis, a closed mouth (asterisk), a pharynx, and an apical tuft (arrowhead) compared to a homozygous *APC* mutant (F) and an AZK treated embryo (G). The latter two (F, G), show a flattened morphology, no pharynx and a secondarily widely open mouth (asterisk).

(**H**) In situ hybridization analysis of *APC* and known "saturating" and "window" genes at the late gastrula stage. Oral views are shown below the corresponding lateral views. Asterisks on lateral views indicate the blastopore. The genotype of the embryo is shown in the upper right corner of each photo. +/+ wild type; +/- heterozygous *APC* mutant; -/- homozygous *APC* mutant. *APC* is expressed in the endoderm, the forming pharynx and in a shallow aboral-to-oral gradient in the ectoderm. *Axin* and *Tcf*, in contrast, are expressed in an oral-to-aboral gradient with a second area of stronger expression at the aboral boundary of the midbody domain. *APC* behaves as a saturating gene in the APC mutant (just as *Axin*, *Tcf*, *Wnt3* and *WntA* – as previously described for AZK treatments (*4*)). *Wnt1* and *Wnt2* behave as window genes in the *APC* mutants and upon AZK treatment (*4*). The only discrepancy in the expression behavior was observed in the case of *Wnt4*, which, for a yet unknown reason, behaves as a saturating gene in the *APC* mutant, but as a “window” gene in the AZK(*4*) (red frame). Lateral views (oral to the left) and oral views are shown.

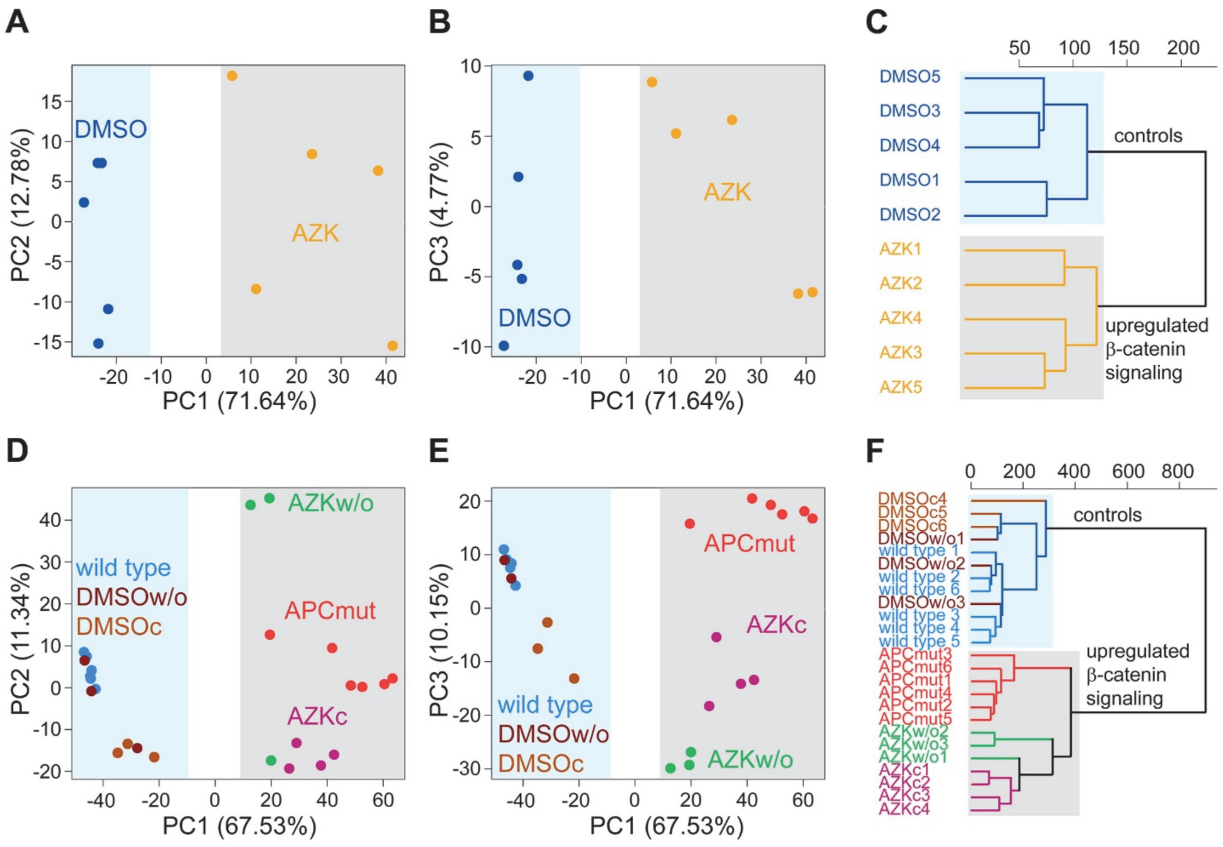


**Supplementary Figure 2 Comparison of transcriptomes of different treatments.**
(**A** to **C**) Principle component analysis and a cluster dendrogram clearly separate the transcriptomes of the biological replicates of the AZK treated (grey background) and DMSO treated (blue background) 1dpf embryos.

(**D** to **F**) Principle component analysis and a cluster dendrogram clearly separate the transcriptomes of the biological replicates of the AZK treated embryos and APC mutants (grey background) versus DMSO treated and untreated wild type (blue background) 3dpf embryos. AZKc – continuous AZK treatment for 3d, AZKw/o – 1 day of AZK treatment followed by a washout for 2 days.

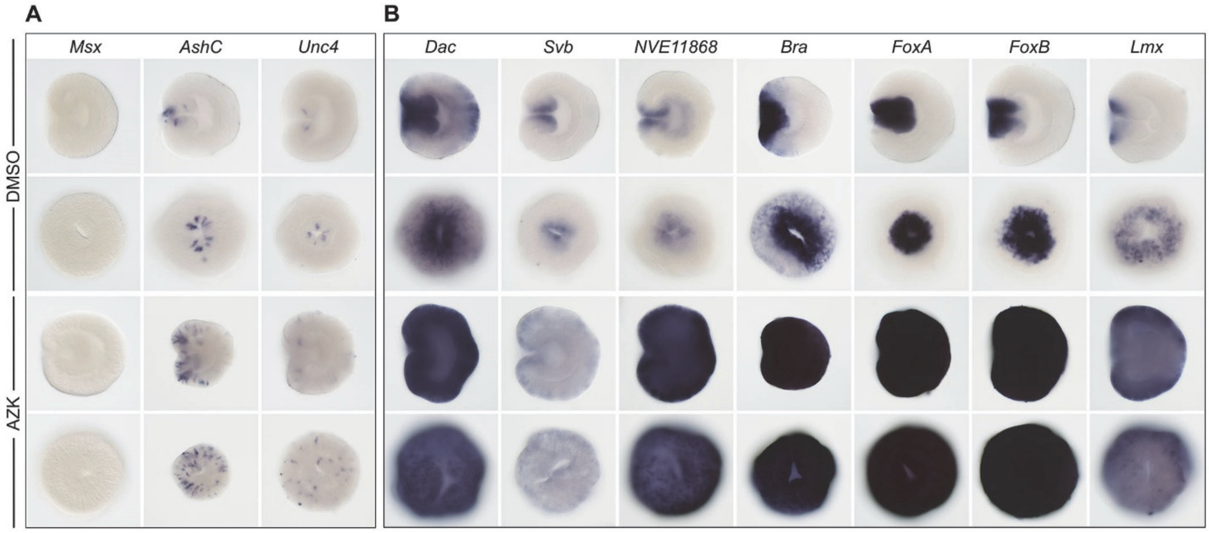


Fig. S3

**Supplementary Figure 3 Expression of ten repressor X candidates in DMSO and AZK.** *Msx* is not detectable at 1 dpf, *AshC* and *Unc4* increase their expression in AZK but are expressed in individual cells. The seven remaining genes display a typical saturating phenotype. Lateral views (oral to the left) and oral views are shown.

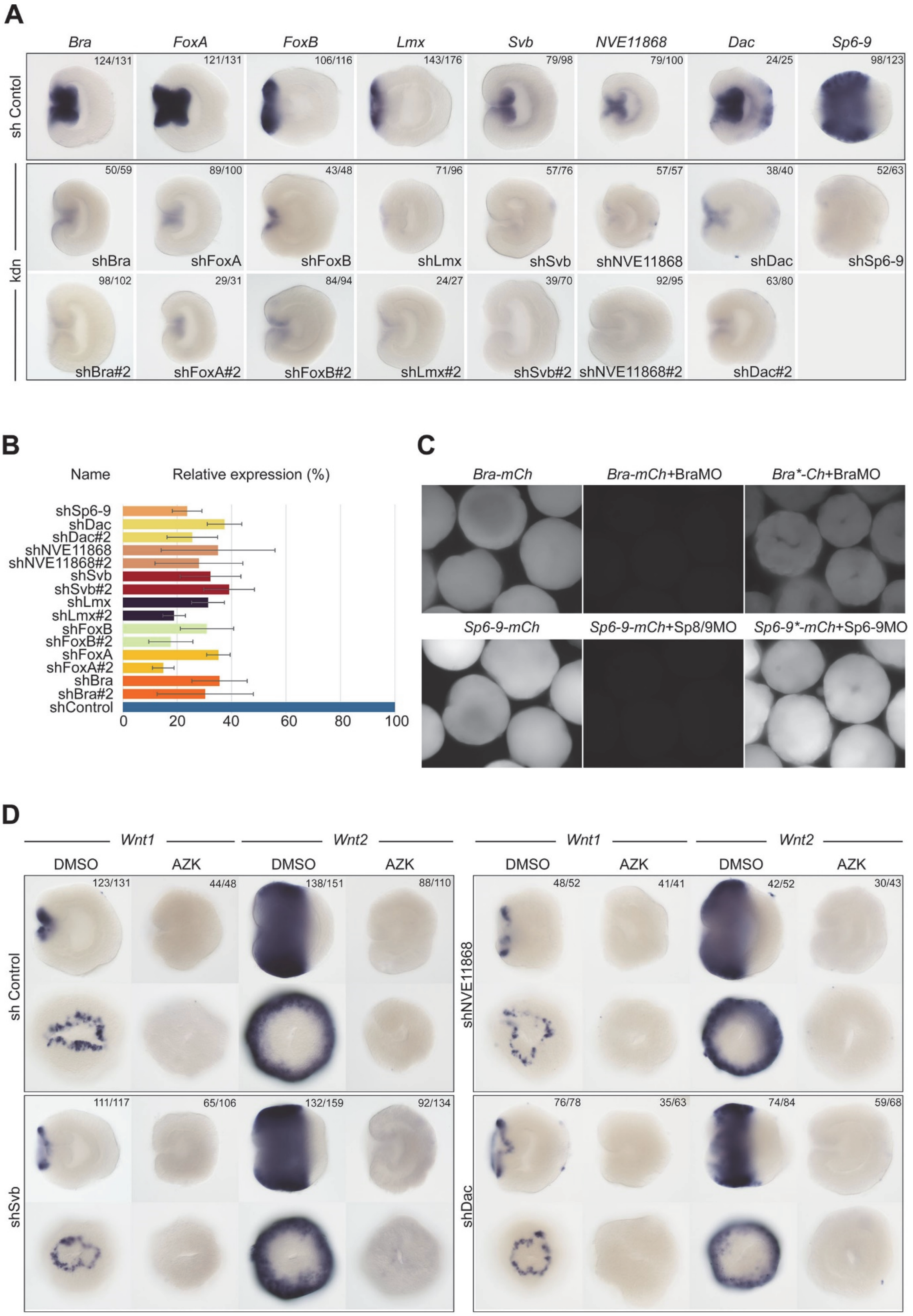
A
Bra
FoxA
FoxB
Lmx
Svb
NVE11868
Dac
Sp6-9
sh Contol
124/131
121/131
106/116
143/176
79/98
79/100
24/25
98/123
kdn
50/59
89/100
43/48
71/96
57/76
57/57
38/40
52/63
shBra
shFoxA
shFoxB
shLmx
shSvb
shNVE11868
shDac
shSp6-9
98/102
29/31
84/94
24/27
39/70
92/95
63/80
shBra#2
shFoxA#2
shFoxB#2
shLmx#2
shSvb#2
shNVE11868#2
shDac#2
B
Name
Relative expression (%)
shSp6-9
shDac
shDac#2
shNVE11868
shNVE11868#2
shSvb
shSvb#2
shLmx
shLmx#2
shFoxB
shFoxB#2
shFoxA
shFoxA#2
shBra
shBra#2
shControl
0
20
40
60
80
100
C
Bra-mCh
Bra-mCh+BraMO
Bra*-Ch+BraMO
Sp6-9-mCh
Sp6-9-mCh+Sp8/9MO
Sp6-9*-mCh+Sp6-9MO
D
Wnt1
Wnt2
DMSO
AZK
sh Control
123/131
44/48
138/151
88/110
shNVE11868
48/52
41/41
42/52
30/43
shSvb
111/117
65/106
132/159
92/134
shDac
76/78
35/63
74/84
59/68

Fig. S4

**Supplementary Figure 4 Testing the efficiency of the shRNAs and morpholinos, and identification of the three candidates not fulfilling the last repressor X criterion.**

For each gene (except *Sp6-9*, for which only one shRNA out of six proved to be effective), two shRNAs have been selected. For *Bra* and *Sp6-9*, translation blocking morpholinos were used as alternative means of knockdown (kdn).

(**A**) In situ hybridization shows reduction in the staining intensity upon shRNA mediated knockdown of the repressor X candidates. Lateral views (oral to the left) are shown.

(**B**) qPCR quantification of the knockdown efficiency for all the shRNAs used in this study. For each shRNA, qPCR was performed on biological triplicates or quadruplicates. The data were normalized to GAPDH expression, and the expression is shown in percent relative to the shControl condition (set to 100%).

(**C**) When co-injected, BraMO and Sp6-9MO bind mRNAs containing their recognition sequence fused to the mCherry coding sequence (*Bra-mCh* and *Sp6-9-mCh*) and suppress their translation. In contrast, no repression of translation is observed when morpholinos are coinjected with mRNAs containing their 5-mismatch recognition sequences fused to the mCherry coding sequence (*Bra*-mCh* and *Sp6-9*-mCh*).

(**D**) *Wnt1* and *Wnt2* are expressed normally in DMSO and are not de-repressed in AZK upon *Svb*, *NVE11868* and *Dac* knockdown. Lateral views (oral to the left) and oral views are shown.

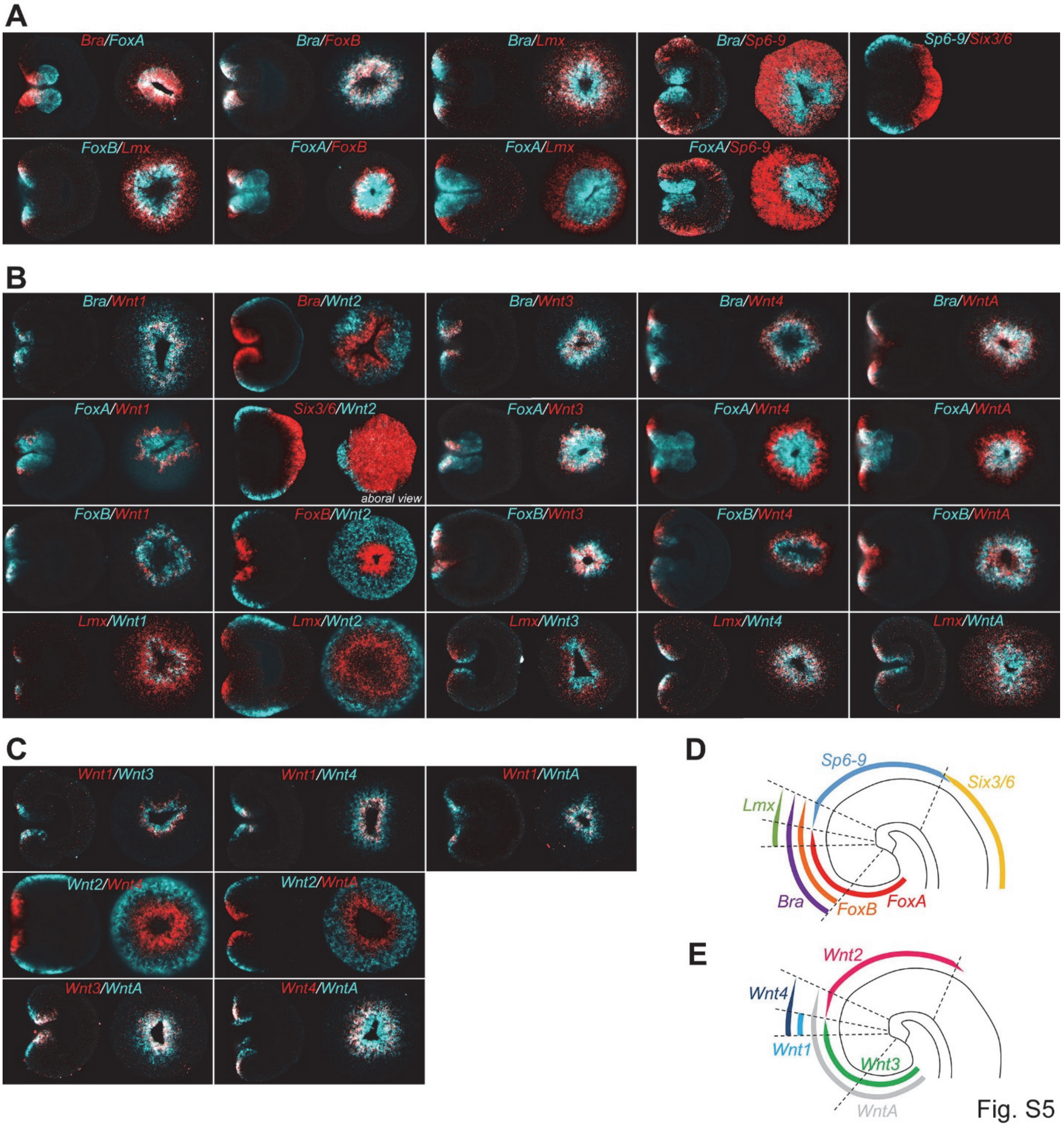


**Supplementary Figure 5 Double FISH analysis of the expression domains of the four main repressor X candidates, oral *Wnt* genes, midbody markers *Wnt2* and *Sp6-9*, and aboral marker *Six3/6*.**

(**A**) FISH analysis of the expression domains of the transcription factors in relation to each other.

(**B**) FISH analysis of the expression domains of the transcription factors in relation to the expression domains of Wnt genes.

(**C**) FISH analysis of the expression domains of the Wnt genes in relation to each other.

(**D**) Schematic representation of the expression boundaries of the transcription factors.

(**E**) Schematic representation of the expression boundaries of the Wnt genes.

Lateral views (oral to the left) and oral views are shown unless specified otherwise.

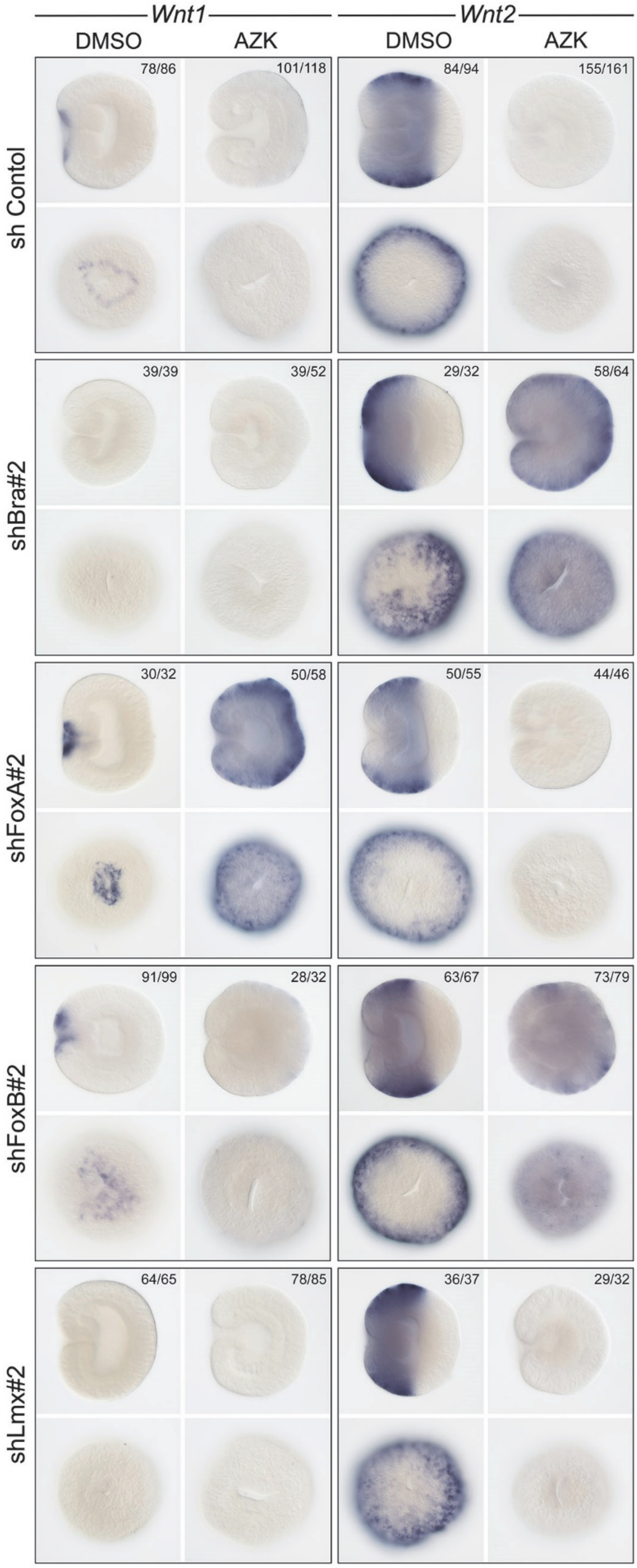

Wnt1
Wnt2
DMSO
AZK
DMSO
AZK
sh Contol
78/86
101/118
84/94
155/161
shBra#2
39/39
39/52
29/32
58/64
shFoxA#2
30/32
50/58
50/55
44/46
shFoxB#2
91/99
28/32
63/67
73/79
shLmx#2
64/65
78/85
36/37
29/32


Fig. S6

**Supplementary Figure 6 Effects of second shRNAs on *Wnt1* and *Wnt2* expression are similar to the effects of the first shRNAs.**

Compare to Fig. 2 in the main text. Lateral views (oral to the left) and oral views are shown.

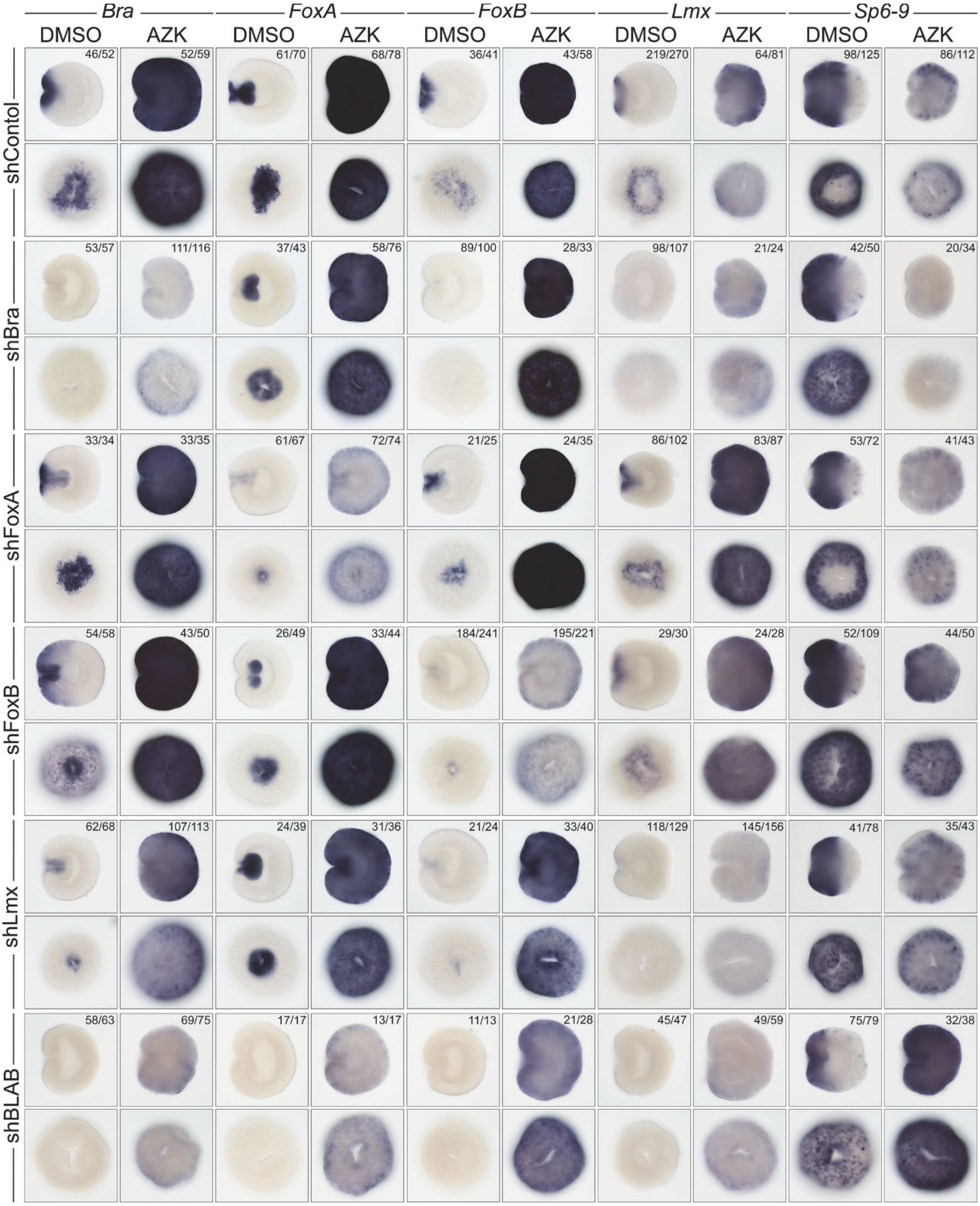
Bra
FoxA
FoxB
Lmx
Sp6-9
DMSO AZK DMSO AZK DMSO AZK DMSO AZK DMSO AZK
shContol
46/52 52/59 61/70 68/78 36/41 43/58 219/270 64/81 98/125 86/112
shBra
53/57 111/116 37/43 58/76 89/100 28/33 98/107 21/24 42/50 20/34
shFoxA
33/34 33/35 61/67 72/74 21/25 24/35 86/102 83/87 53/72 41/43
shFoxB
54/58 43/50 26/49 33/44 184/241 195/221 29/30 24/28 52/109 44/50
shLmx
62/68 107/113 24/39 31/36 21/24 33/40 118/129 145/156 41/78 35/43
shBLAB
58/63 69/75 17/17 13/17 11/13 21/28 45/47 49/59 75/79 32/38

Fig. S7

**Supplementary Figure 7 Effects of the individual knockdowns and of the simultaneous knockdown of all four repressor X candidates on their own expression and on the expression of *Sp6-9*.**

Lateral views (oral to the left) and oral views are shown. See discussion in the main and supplementary text.

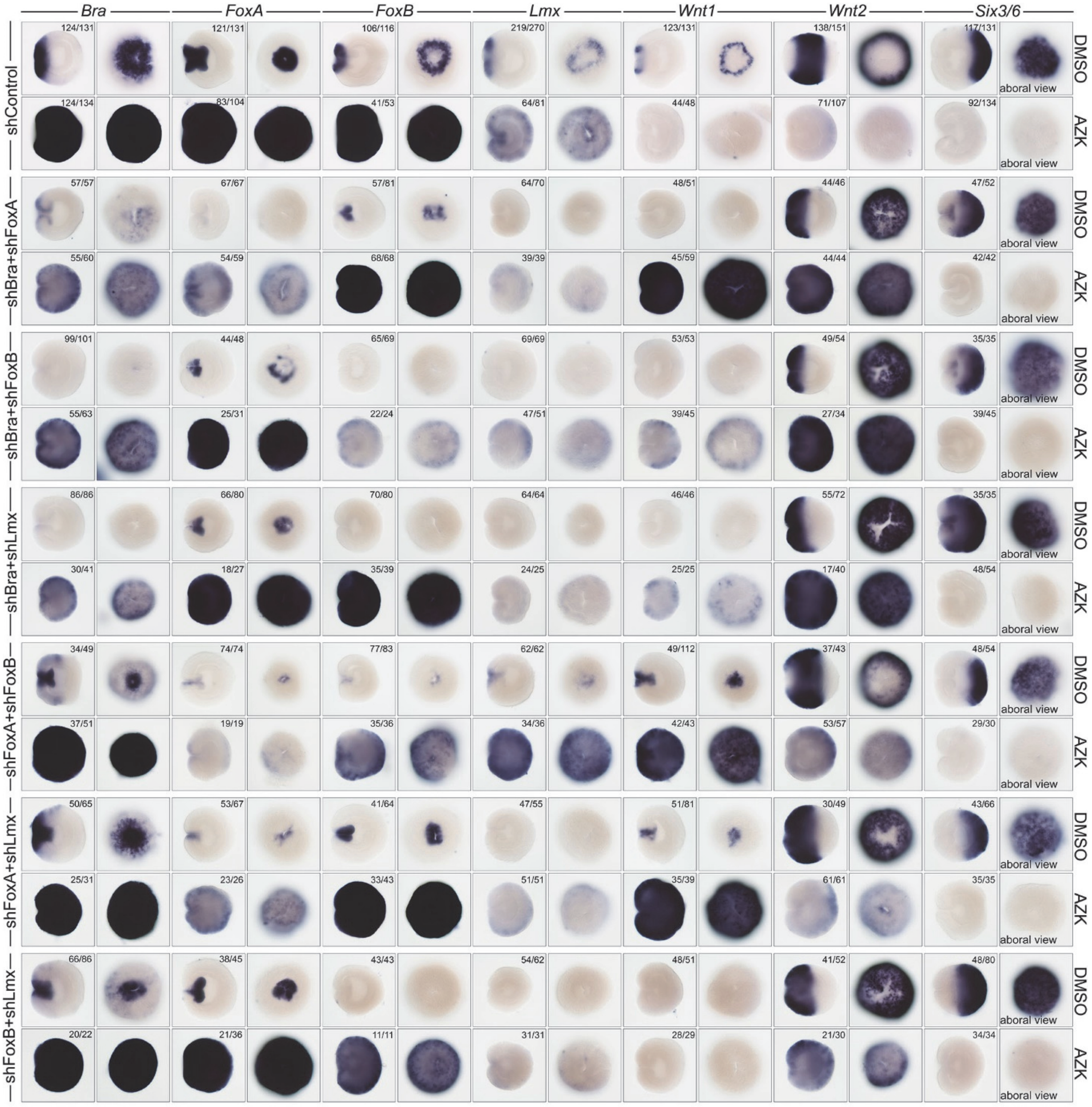


Fig. S8

**Supplementary Figure 8 Effects of the double knockdowns of all possible combinations of the four repressor X candidates on their own expression and on the expression of *Wnt1*, *Wnt2*, and of the aboral marker *Six3/6*.**

Lateral views (oral to the left) and oral views are shown. See discussion in the main and supplementary text. Note oral expansion of Six3/6 upon knockdowns with shRNA combinations containing *Lmx* and, especially, *Bra*. *Bra* knockdown also leads to the expression of *Six3/6* at the bottom of the pharynx.

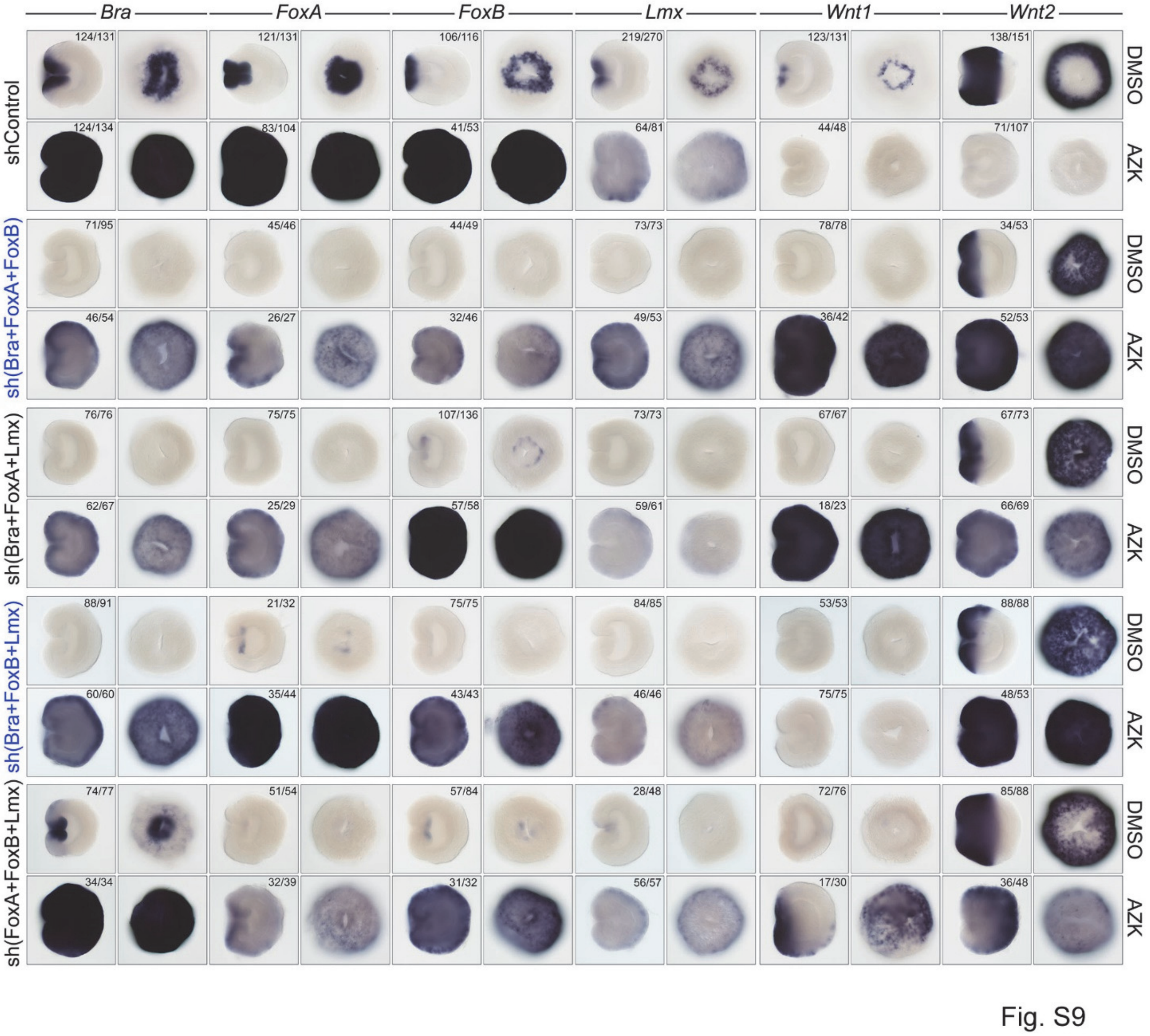


**Supplementary Figure 9 Effects of the triple knockdowns of all possible combinations of the four repressor X candidates on their own expression and on the expression of *Wnt1*, and *Wnt2*.**

Lateral views (oral to the left) and oral views are shown. See discussion in the main and supplementary text.

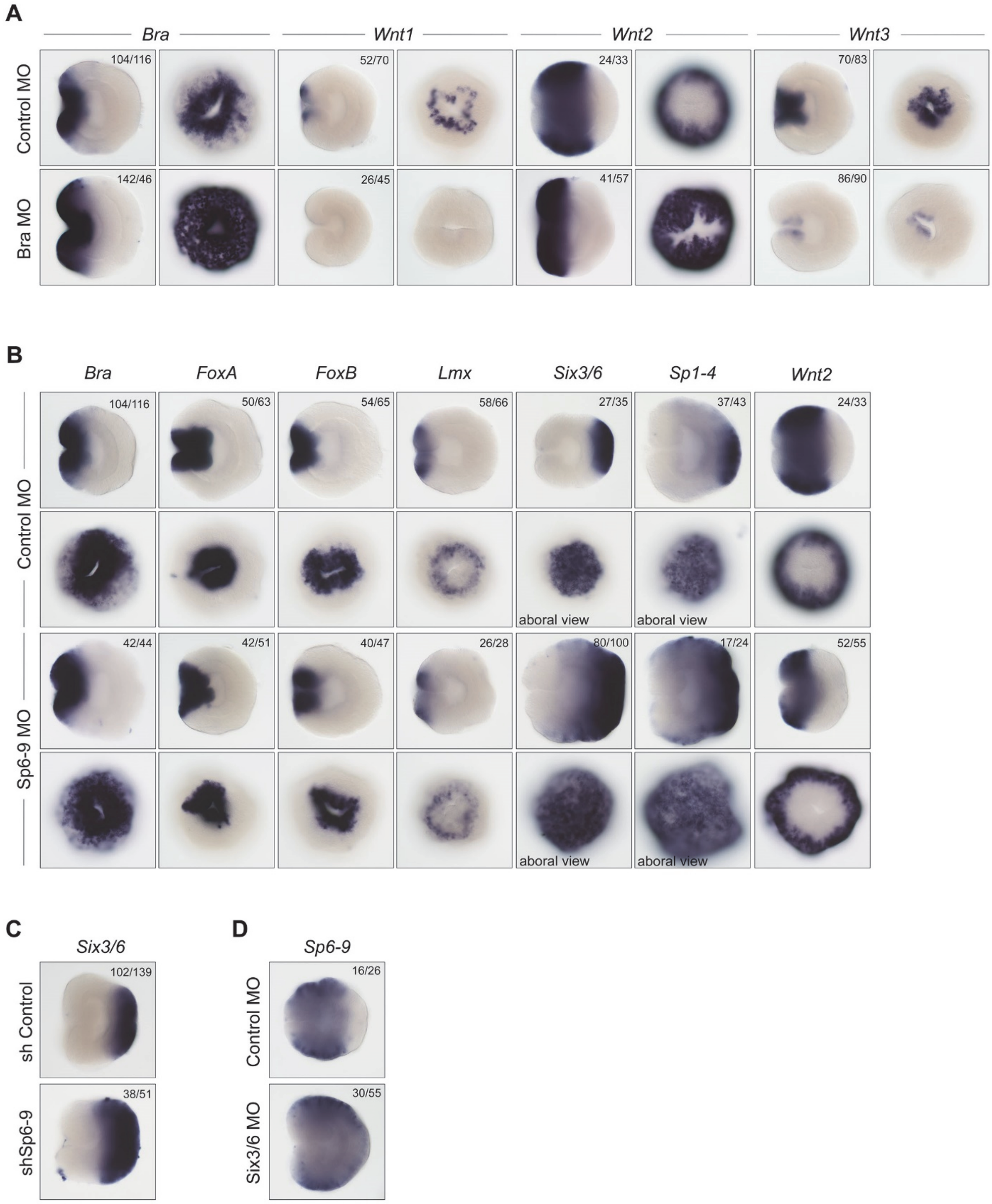


**Supplementary Figure 10 Morpholino experiments**

(**A**) Injection of BraMO has the same effect on *Wnt1*, *Wnt2* and *Wnt3* expression as shBra. In contrast, *Bra* expression is clearly upregulated upon BraMO injection suggesting a negative feedback loop.

(**B**) Sp6-9MO does not affect *Bra*, *FoxA*, *FoxB* and *Lmx* expression. *Wnt2* ring becomes narrower, while *Six3/6* and *Sp1-4* expression domains expand orally.

(**C**) shSp6-9 affects *Six3/6* expression in the same way as the Sp6-9MO.

(**D**) Six3/6MO injection leads to aboral expansion of *Sp6-9*.

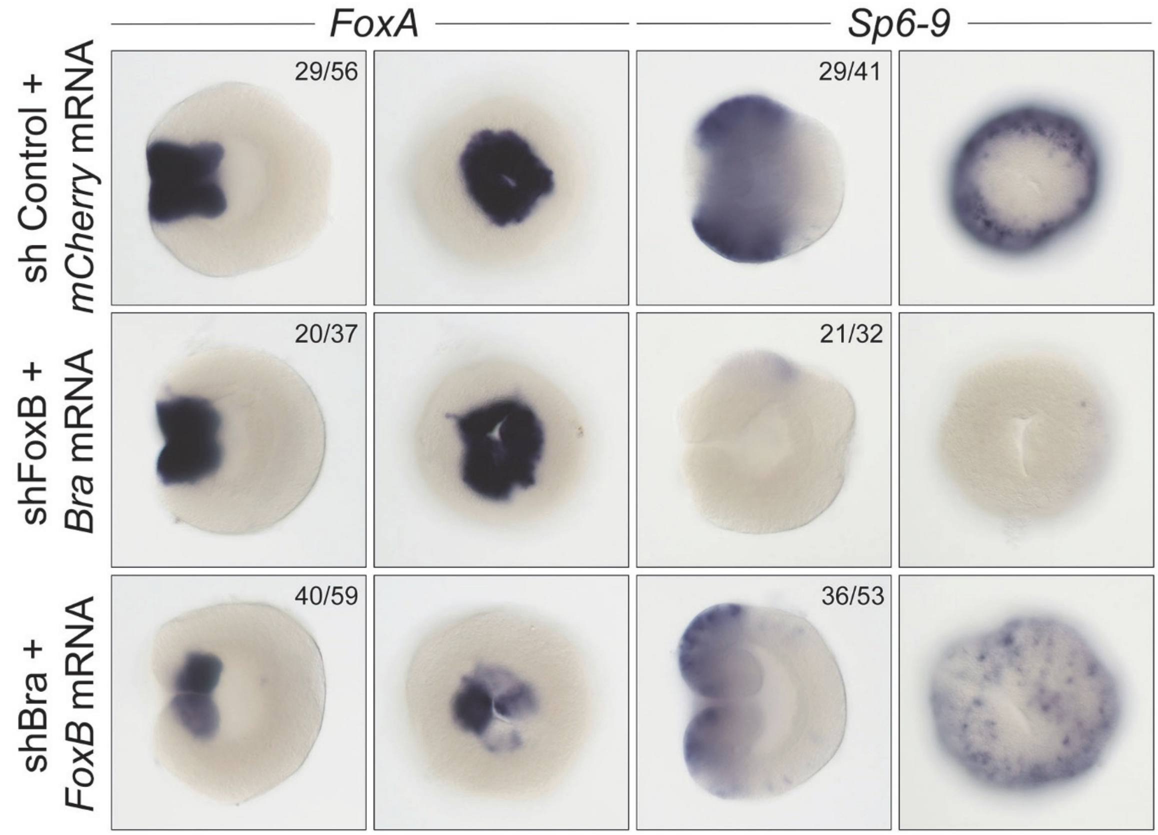


Fig. S11

**Supplementary Figure 11 Rescue experiments with *Bra* and *FoxB***

Ubiquitous *Bra* expression compensates for the lack of FoxB and rescues *FoxA* in its normal domain without causing ectopic overexpression. In contrast, *FoxB* mRNA does not rescue the shBra effect on *FoxA* expression. *Bra* overexpression abolishes *Sp6-9* irrespective of the lack of FoxB, while *FoxB* overexpression makes the shBra effect on *Sp6-9* milder without alleviating it completely (weaker *Sp6-9* expression with a milder oral expansion in comparison to shBra alone). Lateral views (oral to the left) and oral views are shown. See fig. S7 for the shBra and shFoxB knockdown phenotypes.

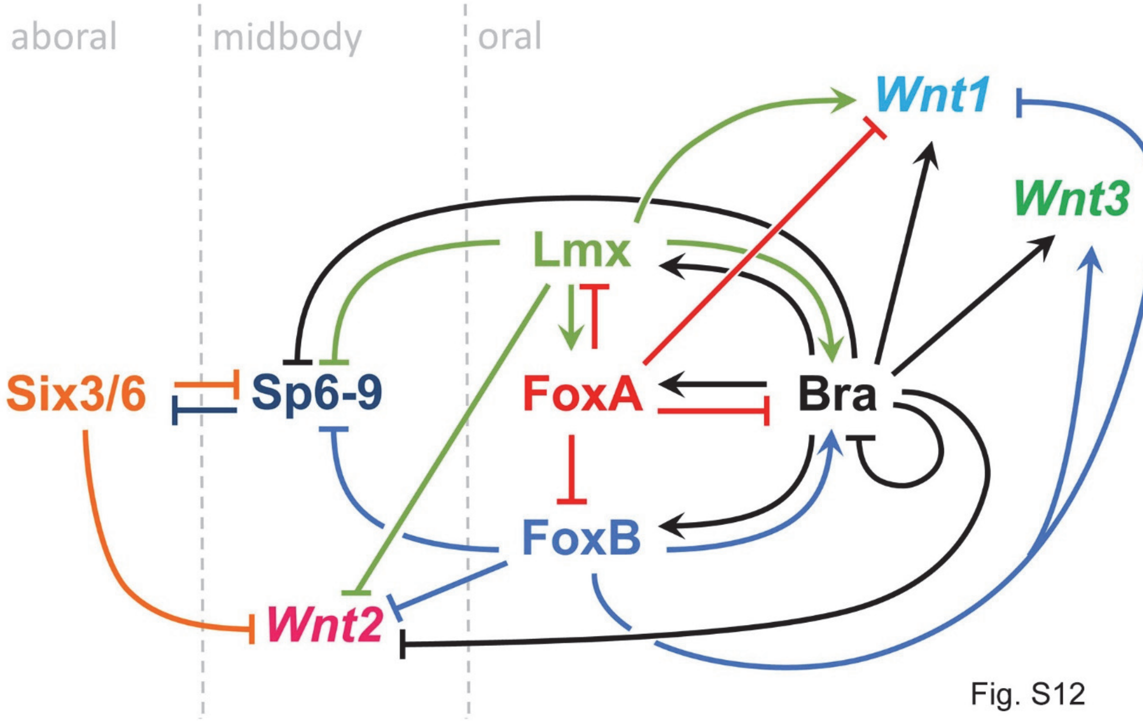


Fig. S12

**Supplementary Figure 12 Putative topology of the GRN of the β-catenin dependent oral-aboral patterning**

The GRN explains why the midbody domain does not expand into the oral, and why the aboral domain does not expand into the midbody. It does not explain, however, why the oral domain does not expand aborally.

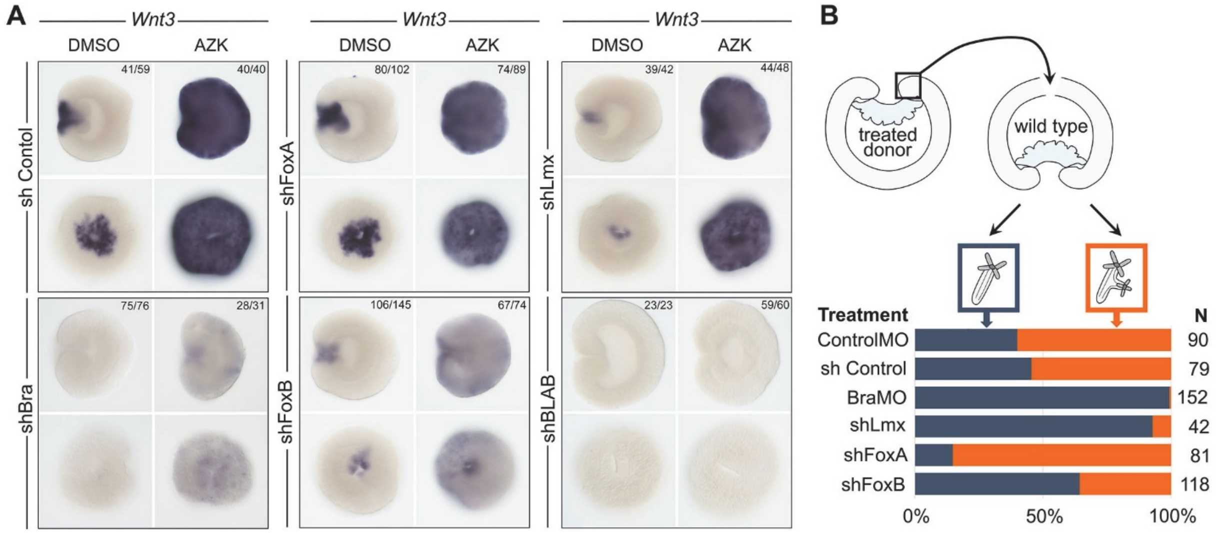


Fig. S13

**Supplementary Figure 13 The effect of the knockdown of repressor X candidates on the inductive capacity of the blastopore lip**

(**A**) Effect of the repressor X candidate knockdown on *Wnt3* expression.

(**B**) The ectopic axis induction capacity of the blastopore lip fragment increases drastically if *FoxA* is knocked down in the donor and sinks if *Bra* or *Lmx* in the donor are suppressed.

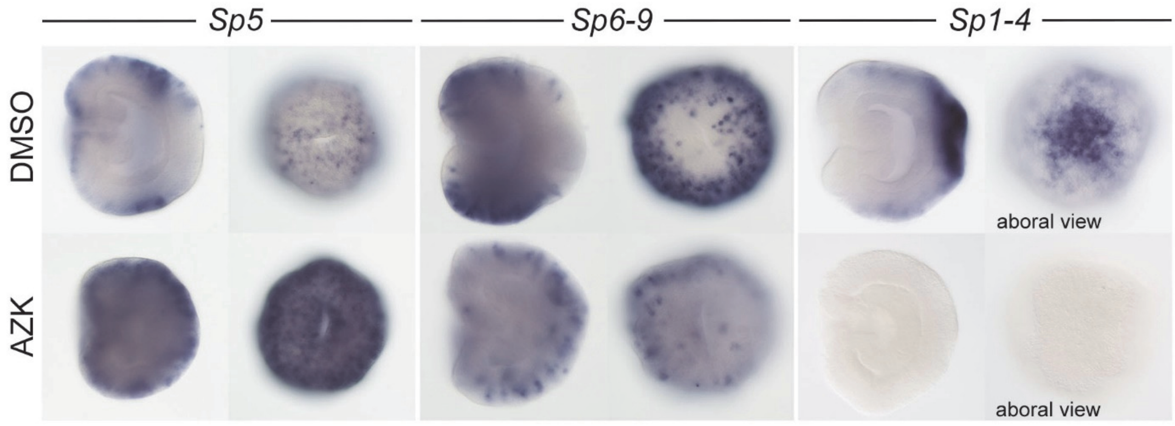


Fig. S14

**Supplementary Figure 14 Expression of *Nematostella Sp* genes in DMSO and AZK**

*Sp5* is a “saturating” gene upregulated in all our 3d treatments according to RNASeq, and also at 1d according to in situ hybridization. *Sp6-9* and, likely, *Sp1-4* are “window” genes.

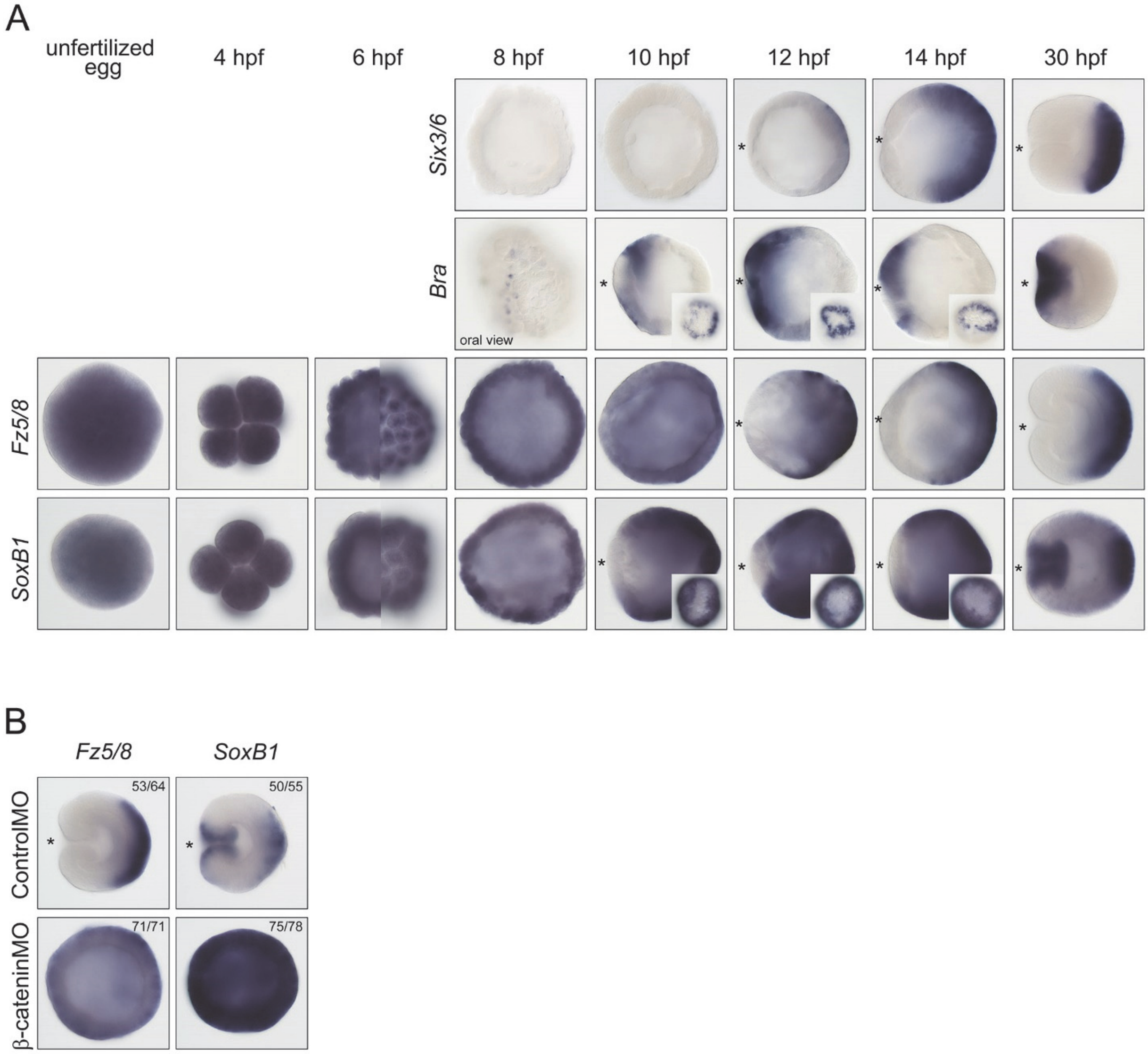


**Supplementary Figure 15 Early expression of *Nematostella Six3/6*, *Bra*, *Fz5/8* and *SoxB1***
(**A**) *Six3/6* is detectable in the aboral portion of the embryo from 12 hpf on. *Bra* becomes detectable in a group of cells on the future oral side of the embryo as early as 8 hpf, and by 10 hpf it forms a ring around the future preendodermal plate. *Fz5/8* is a maternally deposited transcript. *Fz5/8* expression shifts to the future aboral side by 12 hpf. *SoxB1* is also a maternally deposited transcript. The loss of *SoxB1* staining in the future endodermal territory occurs simultaneously with the formation of the *Bra* ring, and is likely regulated by the same mechanism. By gastrula stage, *SoxB1* is expressed in the blastopore lip and aborally (see also Suppl. Fig. 16A). On all lateral views, on which the O-A axis is discernible, the oral end is marked with an asterisk. Inset images of 10, 12 and 14 hpf embryos stained for *Bra* and *SoxB1* show the lack of expression in

the putative preendodermal plate on embryos orientated with their oral ends facing the viewer. 6hpf images of *Fz5/8* and *SoxB1* expression show the optical midsection (left) and the surface view (right) of the same embryos.

(**B**) *Fz5/8* and *SoxB1* expression remains ubiquitous in the β-catenin morphants. Lateral views of the 30 hpf gastrulae, oral ends are marked with an asterisk.

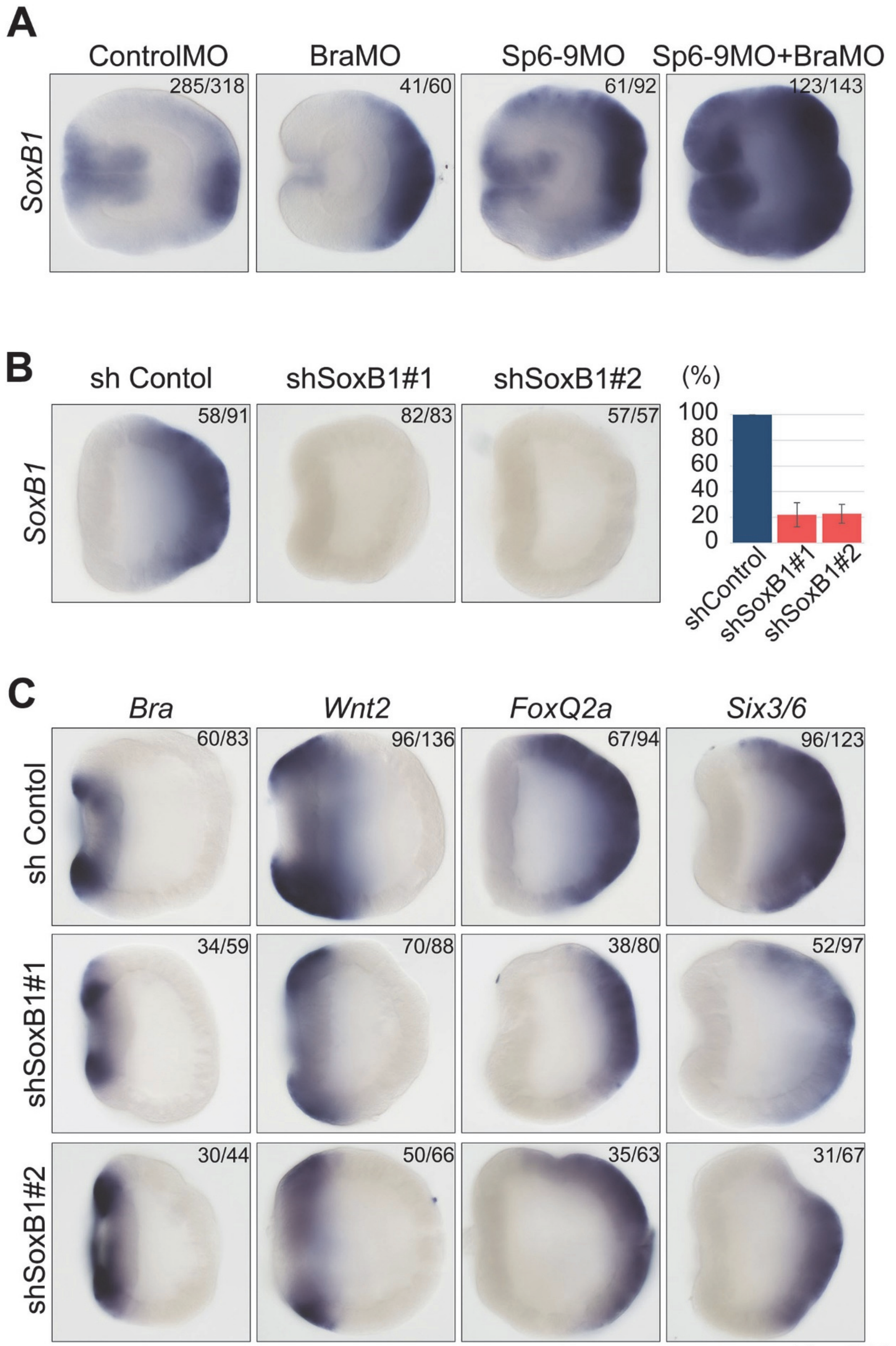
A
ControlMO
BraMO
Sp6-9MO
Sp6-9MO+BraMO
SoxB1
285/318
41/60
61/92
123/143
B
sh Contol
shSoxB1#1
shSoxB1#2
SoxB1
58/91
82/83
57/57
(%)
100
80
60
40
20
0
shControl
shSoxB1#1
shSoxB1#2
C
Bra
Wnt2
FoxQ2a
Six3/6
sh Contol
shSoxB1#1
shSoxB1#2
60/83
96/136
67/94
96/123
34/59
70/88
38/80
52/97
30/44
50/66
35/63
31/67

Fig. S16

**Supplementary Figure 16 *SoxB1* is repressed in the midbody domain and does not seem to be the critical regulator of *Six3/6* and FoxQ2a expression**

(**A**) *SoxB1* expression upon *Bra* knockdown appears weaker in the oral domain and expanded in the aboral domain, which is likely due to the oral shift of the *Sp6-9* expression. *Sp6-9* knockdown significantly expands *SoxB1* expression fusing the oral and aboral expression domains. Simultaneous knockdown of *Bra* and *Sp6-9* makes this effect even more pronounced consistent with the general aboralization of the embryo.

(**B**) *SoxB1* is efficiently knocked down by two independent shRNAs, as demonstrated by in situ hybridization and qPCR.

(**C**) *SoxB1* knockdown does not affect the expression of oral and midbody markers *Bra* and *Wnt2*, and appears to slightly reduce the expression of the aboral markers *Six3/6* and *FoxQ2a* at the 18 hpf pregastrula stage.

**Table S1 Short hairpin RNA targets**

| Name | Targeted sequence |
|---|---|
| shControl | GCGAGTTCTTCTACAAGGTGA |
| shBra#2 | GAAGAGATCACGAGTCTAA |
| shBra | GAATCGCACTCAGCTTACT |
| shFoxA#2 | GCAGGTATGCCCATGAATA |
| shFoxA | GCTCAAGAAATCCAAGGACAA |
| shFoxB#2 | GAGAAGACGAGGATGAACT |
| shFoxB | GATTCCCTCTCTTCCTACA |
| shLmx#2 | GTGTCACATTCTCCGTACAT |
| shLmx | GCTTGAGTGTAAGAGTGGT |
| shSvb#2 | GAACCTTAGATCGGAGAGA |
| shSvb | GCTCCGAGAAGAGAATGTT |
| shNVE11868#2 | GAATGACCTTGAGTGAAGA |
| shNVE11868 | GGAGAGAGAGGTAACTGAT |
| shDac#2 | GCAGAACAGCGAGTAACAA |
| shDac | GACTCTACTGAGGAACATA |
| shSp6-9 | GCTTGAGGGATCGACTTCA |
| shSoxB1 | GCAGCACAGTCCTTTAATA |
| shSoxB1#2 | GGATCCTACTCGAACATGT |

See Fig. S 4 and S16 for the knockdown efficiency estimation by qPCR and in situ hybridization

**Table S2 Morpholino sequences**

| Name | Morpholino sequence | Reference, DOI |
|---|---|---|
| BraMO | TCGTCCGAGTGCATGTCCGACTATG | new |
| Sp6-9MO | TCTAGTAGTTCCTGTGAGTAGACAG | new |
| ControlMO | GATGTGCCTAGGGTACAACAACAAT | Kraus et al., 2016<br>10.1038/ncomms11694 |
| Six3/6MO | GTACTGCCGCACTGCAAGACTTGTC | Sinigaglia et al., 2013<br>10.1371/journal.pbio.1001488 |
| β-cateninMO | TTCTTCGACTTTAAATCCAACTTCA | Leclère et al., 2016<br>10.1242/dev.120931 |

See Suppl. Fig. 4 for the confirmation of the sequence-specific activity of the new MOs.
